# A ‘one-two punch’ therapy strategy to target chemoresistance in estrogen receptor positive breast cancer

**DOI:** 10.1101/2020.03.12.989251

**Authors:** Feng Chi, Jiayi Liu, Samuel W. Brady, Patrick A. Cosgrove, Aritro Nath, Jasmine A. McQuerry, Sumana Majumdar, Philip J. Moos, Jeffrey T. Chang, Michael Kahn, Andrea H. Bild

## Abstract

**Background:** Cancer cell phenotypes evolve over the course of a tumor’s treatment. The phenotypes that emerge and disappear over time will be specific to each drug regimen and type of cancer. Chemotherapy remains one of the most common and effective treatments for metastatic breast cancer patients; however, resistance to chemotherapy inevitably emerges. Cancer chemotherapy treatment regimens are not designed to target emerging chemo-resistance, despite its clear importance in progressive cancer. This study focuses on finding sequential treatment strategies that target acquired chemo-resistant states and optimize response to chemotherapy.

**Methods:** In this study, we used heterogeneous tumor samples from patients to identify subclones resistant to chemotherapy. Using flow cytometry for stem cell markers and DNA sequencing to define subclonal population changes, we measured the enrichment of cancer stem cell-like (CSL) phenotypes in subclones that survive chemotherapy. We then analyzed breast cancer patient tumor organoids and cell line acquisition of CSL traits following chemotherapy, as well as the ability of different drugs to reverse acquired resistance, using flow cytometry, mammosphere assays, and single cell RNA-sequencing analysis.

**Results:** We show that in progressive estrogen receptor positive (ER+) metastatic breast cancer patients, resistant tumor subclones that emerge following chemotherapy have increased CSL abundance. Further, in vitro organoid growth of ER+ patient cancer cells also shows that chemotherapy treatment leads to increased abundance of ALDH+/CD44+ CSL cells. Chemotherapy induced CSL abundance is blocked by treatment with a pan-HDAC inhibitor, belinostat. Further, belinostat treatment diminished both mammosphere formation and size following chemotherapy, also indicating a decrease in progenitor CSL traits. HDAC inhibitors specific to class IIa (HDAC4, HDAC5) and IIb (HDAC6) were shown to primarily reverse the chemo-resistant CSL state. Single-cell RNA sequencing analysis with patient samples showed that HDAC targets and MYC signaling were promoted by chemotherapy and inhibited upon HDAC inhibitor treatment.

**Conclusion:** These findings indicate that HDAC inhibition can block chemotherapy-induced drug resistant phenotypes with ‘one-two punch’ strategy in refractory breast cancer cells.

## Background

Breast cancer is the most frequently diagnosed and second leading cause of cancer-related deaths in women [1]. Chemotherapy kills cancer cells by inducing apoptosis through DNA damage and/or mitotic division inhibition [2]. Chemotherapeutics are often standard of care in clinical oncology because of their effectiveness in reducing tumor burden and improving survival [3]. Nevertheless, some patients will recur with metastatic progression, which has a 90% of cancer mortality [4], resulting in a 23% 5-year survival rate for these breast cancer patients [1].

Tumors are composed of heterogeneous populations of cells, thought to have a hierarchical organization driven by cancer stem cells (CSCs). CSCs are a small therapy-resistant sub-population of cells within tumors that possess the capacity of self-renewal and are capable of promoting a refractory state in patients following drug treatment [5]. These CSCs can initiate a new primary tumor, bear an inherent resistance to chemotherapeutic drugs, and contribute to recurrent disease [6]. Breast CSCs (BCSCs) exhibit a CD44 high/CD24 low phenotype with high ALDH1 expression (ALDH1+) [7]. Tumor cells can also acquire stem cell like characteristics, such as expression of progenitor markers and drug resistance. These cells may represent a de-differentiated state similar to CSCs and reflect a more primitive tumor cell progenitor [8].

Based on the mechanisms of action, chemotherapies could be divided into at least three major groups: antimetabolites; genotoxic agents (eg. doxorubicin serving as alkylating agent, which inhibits DNA topoisomerase II and induces DNA damage and apoptosis; carboplatin serving as intercalating agent, which binds in the grooves in the DNA helix and interfering with polymerase activity during replication/transcription); and mitotic spindle inhibitors (eg. paclitaxel, which disrupts mitosis by affecting the formation/function of spindle microtubule fibers required for chromosome alignment) [9]. Some resistance mechanisms have been described for these chemotherapies. For example, chemoresistance can be acquired by altered membrane transport through ABCB1 (P-gp or MDR1) for doxorubicin and paclitaxel [9], and enhanced DNA repair through increased level of excision repair cross-complementing protein (ERCC1) for carboplatin [10]. Current chemotherapeutic regimens target the bulk of the tumor cells and may benefit from also targeting resistant cells, such as cancer stem cells (CSC) or cancer cells that have stem-like traits such as drug resistance or de-differentiated states [11]. Failure to eliminate these cells can lead to drug resistance, subsequent recurrence and metastasis [12], suggesting that targeting these populations may be necessary to improve outcomes [5,13]. Multiple strategies have been proposed to combat CSCs; however, clinical implementation has remained elusive [7, 14]. A strategy combining conventional cytotoxic chemotherapy and anti-CSC compounds could increase efficacy in reducing the risk of breast cancer relapse and metastasis [7]. The drug-tolerant phenotype within a small subpopulation of cancer cells have been found transiently acquired and reversible, which could be selectively ablated by chromatin-modifying agents, such as HDAC inhibitor, suggesting a potential therapeutic opportunity with ‘one-two punch’ strategy [15].

Single-cell sequencing techniques have been leveraged to identify resistant cancer evolution upon chemotherapy treatment [16-19]. Previously, we performed whole-genome sequencing (WGS) and single-cell RNA-Seq (scRNA-Seq) on four patients’ matched pre-treatment (chemo-sensitive) and post-treatment (chemo-resistant) samples to investigate the mechanisms of acquired chemotherapy resistance in breast cancer. Three of four patients demonstrated increased post-treatment stem cell-like and mesenchymal properties, which may have promoted acquired drug resistance in these patients [19]. Based on this data, we sought to understand how multiple different chemotherapies impact CSL state and determine therapy approaches to prevent emergence of CSL cells. Here, we find that chemotherapies with different mechanisms of action can select for subclones with primitive traits through transcriptional changes. Belinostat, a pan-HDAC inhibitor, was found to prevent this selection and reverse CSL therapy resistance signaling, such as signaling in MYC and dedifferentiation pathways. The specific Class II HDAC inhibitors, LMK-235 and CAY10603, which target to HDAC4/5 and HDAC6 respectively, exhibited the similar reversal effect on chemo-treatment acquired stemness.

In total, this research provides the following unique findings: 1) a novel one-two, chemo-belinostat combination therapy strategy to target chemo-treatment acquired stemness, 2) the ability of belinostat to serve as a common anti-CSL inhibitor following multiple chemotherapies, 3) the specificity of HDAC inhibitors to target the stemness phenotypic shift and not just a change in tumor cell viability and/or apoptosis, 4) the use of patient tumor samples to validate that stemness enrichment in subclonal populations of patients is linked to their survival during chemotherapy, 5) an understanding of the transcriptional changes in single tumor cells upon CSL phenotype reversal, and 6) the role of HDAC class IIa and IIb on chemoresistance through their regulation on primitive traits. Taken together, these findings could help to build a new strategy consisting of sequential chemotherapy and HDAC inhibitor combination treatment to combat refractory breast cancer.

## Methods

### Cells and drugs

Breast cancer cell lines CAMA-1, T47D, MCF7, ZR-75-1, MDA-MB-231, BT549, HCC38, HCC1143, and HCC1395 were all obtained from ATCC. BGB324 was obtained from BerGenBio. The following drugs were all purchased from Selleckchem s including: chemotherapy drugs (Doxorubicin, Carboplatin, and Paclitaxel), AXL inhibitors (BGB324 and TP-0903), Wnt/CBP inhibitor (ICG-001,), Hedgehog inhibitor (vismodegib), CDK8 inhibitor (senexinA), CDK4/6 inhibitor (ribociclib and abemaciclib), FGFR inhibitor (AZD4547), EGFR inhibitors (afatinib and gefitinib), ALDH inhibitor (disulfiram, DSF), pan-HDAC inhibitor belinostat, Class I HDAC inhibitor entinostat, and specific HDAC inhibitors SantacruzanateA (HDAC2i), RGFP966 (HDAC3i), PCI-34051 (HDAC8i), LMK-235 (HDAC4/5i), CAY10603 (HDAC6i).

### Sample collection

Primary breast cancer samples were obtained from malignant pleural effusions or ascites. Patients #1-4, #7-8, #11 are from pleural effusions, patient #5-6, #9-10, #12 are from ascites. After fluid drainage, cells were pelleted at 300 × g for 5 min, and resuspended in TAC buffer (17mM Tris, pH 7.4, and 135mM NH4Cl), followed with incubation at 37 °C for 5 min and centrifugation. After repeating these steps several times until red blood cells were completely depleted from pellet, the remaining cells were washed with PBS twice and frozen in 90% FBS with 10% dimethyl sulfoxide (DMSO).

### Treatment of patient cultures/breast cancer cell lines with chemotherapy and CSL inhibitors for FACS analysis

For patient samples used for FACS analysis and single cell RNA sequencing, we thawed frozen viable breast cancer pleural effusions or ascites from patients and depleted dead cells with EasySep™ Dead Cell Removal (Annexin V) Kit (STEMCELL Technologies) and CD45+ cells with EasySep™ Human CD45 Depletion Kit II (STEMCELL Technologies) according to manufacturer protocols. These purified cancer cells were then resuspended in Renaissance Essential Tumor Medium (RETM, Cellaria) containing 5% FBS, RETM supplement, antibiotic/antimycotic (anti-anti), and 20 ng/mL cholera toxin. For chemotherapy dose response evaluation, patient cells were counted and plated at 10,000 per well in 100 µL RETM with 96-well spheroid plates with black walls (Corning 4520). Breast cancer cell lines were counted and plated at 40,000 per well in 100 µL culture medium (RPMI contain 10% FBS, 1% antibiotic-antimycotic for T47D, ZR-75-1, MDA-MB-231, BT549, HCC38, HCC1143, and HCC1395, DMEM containing 10% FBS, 1% antibiotic-antimycotic for CAMA-1 and MCF7). After 18-24 hours, 100 µL culture medium with drug (2X dose) was added. Cells were harvested after 72 hours for ALDEFLUOR and CD44 staining and FACS. For CSL inhibitor treatment, cells were plated and treated with chemotherapy the same as mentioned above. After 72 hours, the medium was removed, and the organoids were washed with fresh medium once and replaced with 200 µL fresh medium containing CSL inhibitors. The cells were harvested after another 72 hours incubation for ALDEFLUOR and CD44 staining and FACS.

### Flow Cytometry

The organoids were collected by centrifugation at 300g for 5min, and dissociated with trypsin-EDTA for 5min at 37°C. The cells were incubated with ALDEFLUOR staining (STEMCELL TECHNOLOGIES) for 30 minutes at 37°C, and then anti-CD44-PE antibody (1:50 Miltenyi) for 25 minutes on ice in ALDEFLUOR Assay Buffer according to the protocol. The samples with DEAB inhibitor without CD44-PE staining serve as the negative control for ALDEFLUOR. For chemo dose response evaluation and CSL inhibitor screening, each sample takes 200 µL /mL for ALDEFLUOR Assay and 100 µL ALDEFLUOR Assay Buffer for CD44 staining. DAPI (1 µg/mL for final concentration) was added to label the dead cells before FACS test.

### Mammosphere Assay

Mammosphere Assay was performed following the protocol mentioned in Shaw L et al with modification (Shaw et al., 2012). Cells were plated in 96-well spheroid plates with black walls (Corning 4520) at 40,000 per/well in 100µL medium. After 18-24 hours, another 100µL medium with 2x drug was added (final concentration Doxorubicin 0.1µM, Carboplatin 50µM, Paclitaxel 1µM). After 72 hours, the spheroids were harvested from 24 wells, and dissociated with Trypsin for 5min at 37°C. The cells were pelleted and resuspended in 1mL mammosphere media (500mL phenol-free RPMI plus 20mL B27, plus rEGF 20ng/mL). For each treatment, the cells were seeded at 18000/well in 6 well plates (ultralow attachment from Corning #3471) with 2mL media supplied with DMSO or Belinostat 5µM in triplicates. After 14 days in culture, the mammospheres were imaged with a Zeiss Axio Observer 7 microscope from triplicate wells. For the second mammosphere culture, the mammospheres from the same treatment were collected and incubated with Trypsin for 5min at 37°C, followed by 10 passages through a 22 gauge needle for dislocation. The cells were seeded again at 18000/well in 6 well ultralow attachment plate with 2mL media supplied with DMSO or Belinostat 5µM in triplicates as the same conditions in the first mammosphere culture. After another 14 days in culture, the mammospheres were imaged again with a Zeiss Axio Observer 7 microscope from triplicate wells. All the mammospheres were counted using Qupath software (a length greater than 80 µm) for both number and area.

### CNV analysis

CNV analysis of four patients’ pre- and post-treatment WGS or WES data were performed and subclonal CNVs were identified as described previously [19]. Briefly, somatic CNVs between integer copy values were considered to be subclonal, such that a region with average CNV 2.5 pre-treatment and which became 3.0 post-treatment was considered to be a mixture of 50% 2-copy cells and 50% 3-copy cells in the pre-treatment sample. For tumor, the Bulk WGS sequencing coverage to identify subclonal CNAs for Patient #1-4 are 60X, 60X, 40X, and 60X, respectively. For germline, the bulk WGS sequencing coverage for Patient #1-4 are 60X, 60X, 40X, and 40X, respectively. For Selecting copy number changes, certain copy number variants were selected by analyzing subclonal, non-integer copy number variants. Other regions do not contribute to this assessment; for example, please reference the horizontal dotted lines along the integer values (at 1, 2, 3, 4, etc.) in S1D-G, which show that copy number variants at these integer positions are not necessary for these analyses. For Stem WGS coverage (1-10X estimated) for comparing subclonal CNA levels between stem cell populations, Low-pass sequencing coverage was used for the ALDH/CD44 sorting for all four patients, with approximately 4X coverage for each sample. The sequencing coverage for Patient #1-4 are 3.3X, 2.5X, 2.5X, and 3.3X, respectively. As shown in Fig. S1D, the therapy-resistant subclones in Patient#1 demonstrated CNVs on chromosomes 3, 4, and 9, whereas therapy-responsive subclones had CNVs on 12, 14, and 16; Patient#2 had responsive CNVs on chromosomes 7q and 11q (Fig. S1E); Patient#3 had resistant CNVs on chromosomes 6 and 7 and a responsive CNV on chromosome (Fig. S1F); and Patient#4 had a resistant (or at least increasing) CNV on chromosome 4 (Fig. S1G). Note that Patient#4’s chromosome 4 amplification does not appear to have changed after treatment, but in fact, increased in the tumor cell population because the sample has much more normal contamination with only ∼25% tumor purity (Fig. S1F).

### CSL proportion calculation in each subclone

To identify CNVs from CSL (ALDH+/CD44+) vs. non-CSL populations, we thawed frozen viable breast cancer pleural effusions or ascites from patients and depleted normal cells, including white blood cells (CD45+), fibroblasts (CD90+), and mesothelial cells (podoplanin+) using the Miltenyi BioTec quadroMACS system with the following protocol. Four mL of quadroMACS buffer (cold PBS with 0.5% BSA and 2 mM EDTA) was added to cells and cells were spun down at 1000 g for 5 minutes at 4 °C. Cells were resuspended in 1 mL quadroMACS buffer, and 10 µL of biotinylated anti-podoplanin antibody (Biolegend #337015) was added. Cells were incubated 10 minutes on ice and spun down at 300 x g for 5 minutes at 4 °C to wash off excess antibody. Cells were resuspended in 120 µL quadroMACS buffer. To this was then added 26 µL of anti-CD45 microbeads, 26 µL of anti-CD90 microbeads, and 26 µL of anti-biotin microbeads (Miltenyi BioTec). Cells were incubated 15 minutes on ice in the dark, followed by addition of 1 mL quadroMACS buffer and cells were then spun down 300 x g for 5 minutes at 4 °C. Cells were resuspended in 500 µL quadroMACS buffer, then added to pre-washed (with 2 mL quadroMACS buffer) LD column, and elute was collected. One mL quadroMACS buffer was added to column and elute continued to be collected; this was repeated an additional time for a final elute volume of 2.5 mL. Cells were then spun down at 1000 g for 5 minutes at 4 °C. We then stained purified breast cancer cells using the ALDEFLUOR system according to manufacturer’s instructions to assess ALDH activity, followed by staining with a CD44-PE antibody in ALDEFLUOR assay buffer for approximately 30 minutes on ice. Cells were washed in ALDEFLUOR assay buffer followed by FACS to assess the number of cells which were ALDH+/CD44+, ALDH+/CD44-, ALDH-, CD44+, or ALDH-/CD44- (referred to as quadrants hereafter), and to sort cells in order to isolate genomic DNA from each of these four populations. Isolated genomic DNA was then sequenced by Illumina whole-genome sequencing at low coverage (2X to 10X) to infer CNVs. To infer CNVs, reads were aligned to hg19 using BWA using the Speed-Seq suite. Then samtools pileup was run on the resulting aligned bam files, and the coverage in 1 million bins across the genome was calculated and manually centered at absolute copy 2 (diploid). The percent of each subclone in each quadrant could be calculated following the example below. An example analysis is shown in Fig. S1B, where a non-stem population (ALDH-/CD44-) had ∼1.6 copies of a chromosome 3 region (40% of cells with chr3 deletion). Meanwhile, the CSL population (ALDH+/CD44+) had ∼1.3 copies of this region (70% of cells with chr3 deletion). Thus, the CSL population is enriched for cells with the chr3 deletion, and these cells represent the survivor subclone. To calculate the percent of each quadrant (CSL population) in each subclone, the survivor subclone and the non-survivor subclones for CSL content are converted following the example below. The number of cells in each stem quadrant (the number of cells for DNA isolation) was recorded to make this conversion. The conversion could be performed like following example described. Suppose the following numbers from FACS in each quadrant were obtained:

1,000 ALDH+/CD44+ cells (CSL cells)

2,000 ALDH+/CD44-cells

2,000 ALDH-/CD44+ cells

10,000 ALDH-/CD44-cells (double-negative)

15,000 cells total

Then assume there were 20% of cells with a subclonal CNV (dubbed CNV-X) in the double-negative non-stem population, but 50% of cells with CNV-X in the CSL population. Based on this, the absolute number of CSL cells with the CNV-X is 1,000 * 0.5 = 500 cells. The absolute number of double-negative cells with CNV-X would be 10,000 * 0.2 = 2,000 cells. This calculation could be performed for each quadrant/subclonal-CNV pair, to get the absolute number of cells in each CSL quadrant for each subclone. Then we sum the total cell number in each subclone in all four quadrants, and calculate the percentage of cells in that subclone that are in each quadrant, with the sum total just described as the denominator. This results in the CSL proportions of each subclone, as shown in Fig. 1A.

**Fig. 1.**
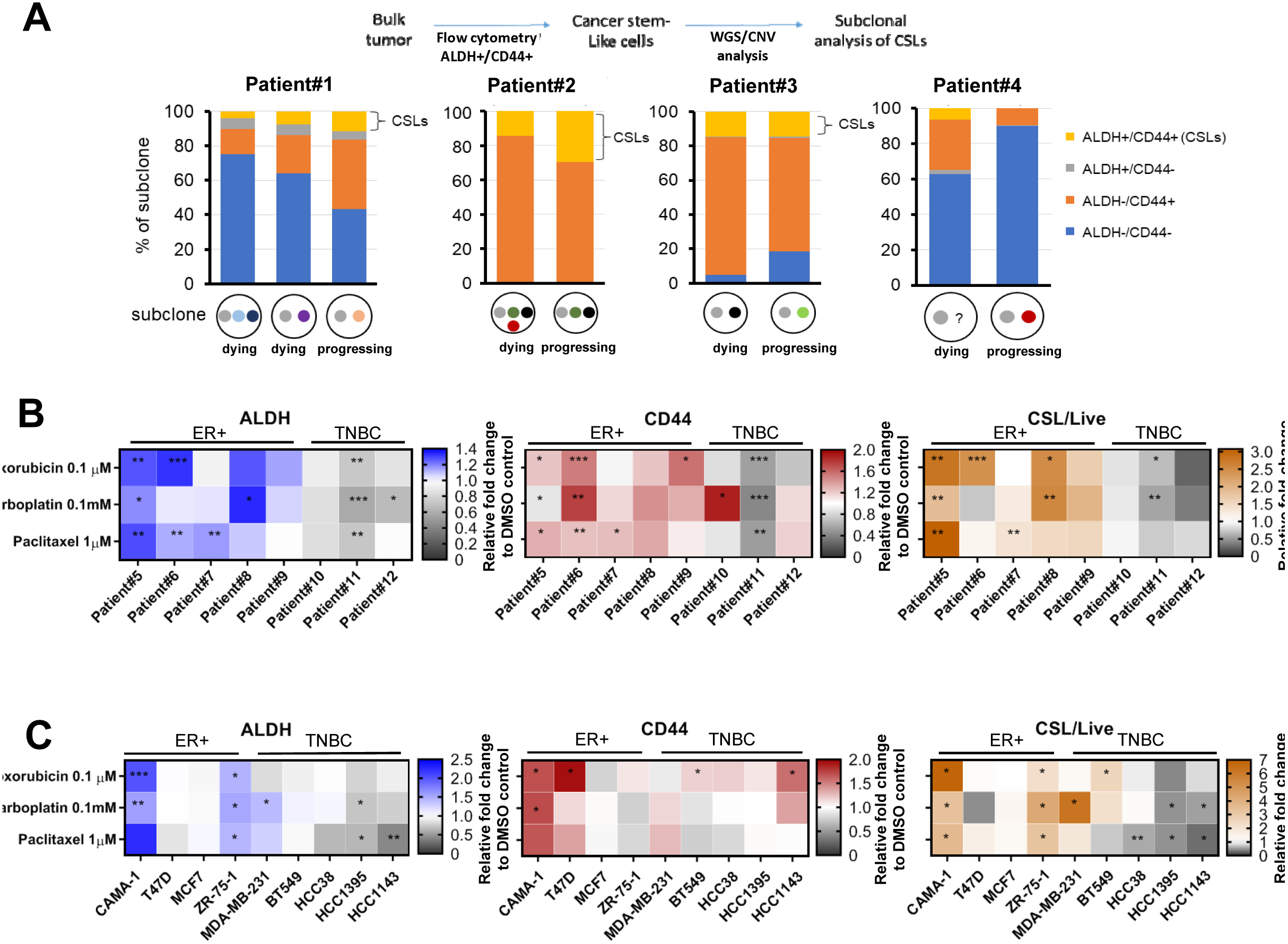
Genetic selection and chemotherapy both enrich CSL subclones. (**A**) The proportion of each subclone with CSL vs. non-CSL populations in four patients. Cells were sorted via FACS using ALDEFLUOR and CD44 staining and DNA was isolated from each population, followed by low-coverage WGS to infer CNV and determine levels of subclonal CNVs. Heat maps for FACS mean values of ALDH, CD44 and CSL cells/Live upon chemotherapy treatment in cultured patient cells (**B**) and breast cancer cell lines (**C**) 3D organoids of patient cells and breast cancer cell lines were incubated with doxorubicin (0.1µM), carboplatin (0.1mM) and paclitaxel (1µM) for 72 hours, followed by ALDEFLUOR/CD44/DAPI staining and FACS to identify the FACS mean values of ALDH and CD44 in live populations and CSL cells/Live. The vehicle controls (DMSO treatment) were set as fold one. All the chemotherapy treatments were expressed as the fold change relative to the control. The mean values of triplicate tests were shown in the heat maps with color indicated as the legends for each treatment. Significance is marked with * for P<0.05, ** for P<0.01, *** for P<0.001.

### Single-cell RNA sample preparation and sequencing analysis

#### Staining of single cell suspensions for scRNA-Seq

The patient cells were isolated, cultured, and treated with chemotherapy +/- belinostat as methods mentioned above for CSL inhibitor screening with FACs analysis. Cell organoids were collected by centrifugation at 300g for 5min, and dissociated with trypsin-EDTA for 5min at 37°C. Dead cells were depleted with EasySepTM Dead Cell Removal (Annexin V) Kit (STEMCELL Technologies, #17899). Cells in suspension were stained with ReadyProbes Cell Viability Imaging Kit (Hoechst 33342 and Propidium Iodide, Thermo Fisher Scientific) by adding 80µL each dye at 500,000 cell/mL in basal DMEM medium and incubated for 20 minutes at 37°C + 5% CO2. After incubation, cells were diluted with equal volume of 1X PBS (no Ca2+/Mg2+, pH 7.4, Thermo Fisher Scientific) then centrifuged for 5 minutes at 300 x g, 4°C. Cells were resuspended to 5.0×10^5^ to 2.0×10^6^ cell/mL in 1X PBS and calculated concentrations on a hemocytometer using trypan blue then diluted to 50,000 cell/mL in 1x PBS + 0.2U/µL Superase. In RNase inhibitor (Thermo Fisher Scientific) + 1X Second Diluent (Takara Bio).

#### Single Cell Dispensing, Imaging, and Candidate Selection

Single-cell RNA-Sequencing (scRNA-Seq) was performed using the ICELL8® Single-Cell System (Takara Bio) using the SMARTer™ ICELL8 3’ DE Kit (Takara Bio). Processing 8 samples at a time, 80µL of each 50,000 cell/mL cell sample, negative control, positive control (K-562 RNA), and fiducial reagent (dye control) were loaded into each well of a 384-well plate (A1 through D2) and dispensed 50nL per well into a 5,184 well SMARTer™ ICELL8® 3’ DE Chip (Takara Bio) using the ICELL8® Multisample NanoDispenser (MSND, Takara Bio). After dispense, chip was sealed with imaging film, and centrifuged at 300 x g, 5 minutes, 4°C. ICELL8® 3’ DE Chip was then imaged on an ICELL8® Imaging System (Takara Bio) featuring an Olympus BX43 upright florescent microscope equipped with an automated stage, camera, DAPI, and Texas Red filters and controlled by CellSelect® Software (Takara Bio) to visually identify and select wells containing one viable single-cell candidate per well (positive Hoechst 33342, negative Propidium Iodide) generating single-cell candidate list for each ICELL8® 3’ DE Chip. After imaging, ICELL8® 3’ DE Chip were stored at -80°C overnight. Each sample was ran twice on separate ICELL8® 3’ DE Chips in order to multiplex samples and minimize chip-to-chip variations.

#### RT-PCR, Library Construction, and Next Generation Sequencing

The ICELL8® 3’ DE Chip is preprinted with a reverse transcription (RT) primer containing a unique molecular identifier (UMI), 11-base unique well barcode, and poly T sequence in each of the 5,184 wells. The ICELL8® 3’ DE Chip containing cells was thawed at room temperature for 10 minutes, then centrifuged for 3 minutes, 3,200 x g, 4°C and maintained on ice. The ICELL8® 3’ DE Chip was placed on the ICELL8® MSND and loaded 50 µL of RT-PCR solution (Takara Bio; 56 µL 5M GC Melt, 24 µL 25mM dNTP Mix, 3µL 1M MgCl2, 9µL 100mM DTT, 62 µL 5X First-Strand Buffer, 33 µL 2X SeqAmp PCR Buffer, 16 µL 10% Triton X-100, 2 µL SMARTer ICELL8 3’ DE Oligo Mix, 29 µL 100U/µL SMARTScribe Reverse Transcriptase, 10µL SeqAMP DNA Polymerase) into four wells (A1 through A4) of a 384-well plate. MSND then dispensed 50 nL to wells containing only live single-cell candidates according to the single-cell candidate file previously generated by the ICELL8® Imaging System. The 3’ DE Chip containing single cell candidates with RT-PCR reagent sealed with PCR Sealing Film and centrifuged 3,200 x g, 3 min, 4°C and then placed in a BioRad T100 equipped with a ICELL8® Chip adapter to perform RT followed by 22 PCR cycles for cDNA amplification. Amplified cDNA from the ICELL8® 3’ DE Chip individual cells was collected and pooled using the ICELL8® Collection Kit (Takara Bio) by centrifugation at 3,200 x g, 10 minutes, 4°C. The cDNA products were concentrated using cDNA Clean & Concentrator-5 Kit (Zymo Research) followed bead cleanup using 0.6X AMPure XP Beads (Beckman Coulter) and eluted in 12 µL DNase/RNase free water.

Libraries were prepared using 1ng of purified cDNA according to the ICELL8® 3’ DE instruction manual (Takara Bio) using the Nextera Primer P5 (ICELL8® 3’ DE Kit, Takara Bio), Nextera XT DNA Library Preparation Kit (Illumina) and i7 Index Primer (Nextera XT Index Kit, Illumina). Unique i7 Indexes were used for each ICELL8® 3’ DE Chip. Tagmented cDNA was amplified by PCR 12 cycles prior to dual-sided bead cleanup using AMPure XP beads, and eluted in 11µL DNase/RNase free water. Library and cDNA quality and profile were confirmed prior to sequencing ran on an Agilent Bioanalyzer using a Bioanalyzer High Sensitivity DNA Analysis kit (Agilent). Prepared library was then sequenced using 150 paired-end cycles on a NovaSeq S4 flow cell (Illumina).

#### Single-cell RNA-Seq Analyses

Libraries were sequenced on an Illumina HiSeq 2500 with 2 × 125 paired-end reads. Raw FASTQ sequencing files were processed by the Cell Ranger Single Cell Software Suite for demultiplexing, barcode assignment, alignment and UMI counting (https://support.10xgenomics.com/single-cell-gene-expression/software/overview/welcome). Samples were aligned to hg19 using the STAR aligner [20]. Count tables were generated with a total of 6196 cells for Patient#5 and Patient#6 and used as input into Seurat v2 [21].

#### Quality control on the cells

Low quality cells were filtered out if the library size was less than 10,000 or the number of expressed genes (counts larger than 0) is less than 500. Furthermore, the cells with more than 30% mitochondrial gene counts were discarded. For Patient#6, 3668 cells were acquired before filtering, 2773 cells were acquired after filtering. For Patient#5, 2528 cells were acquired before filtering, and 1553 cells were acquired after filtering.

#### Dimensionality reduction, clustering and visualization using UMAP

The Bioinformatics ExperT SYstem (BETSY) [22] was applied to cluster and visualize the scRNA-seq data using UMAP. In short, an RNA-Seq count matrix was normalized to log scale with a scale parameter of 10,000. Top 500 highly variable genes were then identified in the normalized dataset. The linear dimensional reduction method PCA was then performed upon these variable genes. The first 10 principal components (PCs) and a resolution of 0.6 were used for Louvain graph-based clustering and two-dimensional visualization with UMAP.

#### Pathway analysis

To identify the pathways reversed by belinostat that are shared by the three chemotherapy groups, we applied the Single Sample Gene Set Enrichment Analysis (ssGSEA) by running the “ssGSEA” function implemented in the GSVA package (v1.22.4) [23] in R (v3.4.3). The gene sets from the hallmark, C2 and C6 collections of the Molecular Signatures Database (v6.2) were used for the ssGSEA. The chemotherapy-belinostat reversal pathways shared among chemotherapy groups were selected with the following steps. Student T-test between chemo+DMSO vs. DMSO+DMSO and chemo+belinostat vs. chemo+DMSO were performed with all ssGSEA enrichment scores for single cells within each chemotherapy group, and all the GSEA pathways with significant adjusted p value (p<0.05) in both T-tests were selected. GSEA pathways with reversal directions in two T-test groups were indicated by the reversal log2 fold changes in the enrichment scores, and filtered with the condition that the pathways must be shared by at least two chemotherapy groups with the same trends. The averaged enrichment scores across all cells in each treatment group were used as the input data to generate heat maps.

### Statistical Analysis

For Single-cell RNA sequencing analysis, the R packages “stats” and “rstatix” were used to perform statistical tests. Student’s t-test (two-tailed) was used to compare the Single Sample Gene Set Enrichment Analysis (ssGSEA) scores between two treatment groups. α level was considered 0.05 for all statistical tests. Error bars for all drug and inhibitor assays represent standard deviation of three replicates from one assay. For all statistical analysis in experiments other than bioinformatics analysis, student’s t-test (two-tailed) was used between groups according to the description in the legends. For all FACS analysis and mammosphere assays, student’s t-tests were performed between treatments vs. controls as indicated in figure legends with triplicate tests. Significance is marked with * for P<0.05, ** for P<0.01, *** for P<0.001.

## Results

### Resistant subclones are enriched in CSL cells

To test how chemotherapy selects for resistant cell traits, we studied CSL characteristics in tumor subclones prior to treatment using four breast cancer patients using copy-number variations (CNV) from WGS [19]. Importantly, while challenging, these studies leverage actual patient tumor phenotype and subclonal evolution during long courses of treatment, for which in vitro work is no substitute. The treatment history for these four ER+ patients and the time points at which analysis was performed are shown in Fig. S1A. To obtain CSL and non-CSL populations, tumor cells collected from each patient were sorted for ALDH and CD44 expression, and cells were isolated to four quadrants (ALDH+/CD44+ CSL cells, ALDH+/CD44-cells, ALDH-/CD44+ cells, and ALDH-/CD44-cells) (Fig. S1B). We defined the CSL population as ALDH+/CD44+ cells [24]. DNA from cells belonging to each of these quadrants was isolated and subjected to low-coverage WGS (2X to 10X) in order to infer copy number variants (CNVs) and determine the prevalence of each subclonal CNV in each cancer stem population (see Fig. S1C). The evolution of subclones is shown in Fig. S1D, E, F, G and described in the Methods. We identified subclone-specific CNVs in each patient at each time point, and then assessed each subclone for the presence of CSL vs. non-CSL populations to determine whether subclones that survive chemotherapy and are also present at time of recurrence had increased numbers of CSL cells. In two of four patients that are progressing on chemotherapy (Patient#1 and Patient#2), the tumor subclone that survives treatment and becomes resistant to chemotherapy has increased CSL abundance compared to other non-resistant subclones (Fig. 1A), suggesting genetic selection can enrich CSL cells in post-treatment resistant subclones in ER+ breast cancer. This result supports our published research showing that patients treated over years of treatments show enrichment in stemness phenotypes [19].

### Chemotherapy-induced feedback increases CSL traits in patient tumor cells

Our previous research demonstrates increased post-treatment stem cell-like properties in three of four patients, including the Patient#1 and Patient#2 used in this work [19]. These findings indicate that the increase in post-treatment CSL cell abundance may be due to chemotherapy-induced feedback. To test this hypothesis, we again use patient cancer cells obtain from malignant pleural effusions or ascites, as these samples may best reflect physiologically relevant phenotypes and signaling changes during drug treatment. These samples were treated with three different chemotherapies of various mechanisms of action (doxorubicin, carboplatin, and paclitaxel) for 72 hours, followed by flow cytometry to measure the CSL abundance using the FACS gating strategy shown in Fig. S2A. Five ER+, PR+, Her2- and three TNBC (triple-negative breast cancer) patient tumor samples were used to test if chemotherapy could induce CSL cells. Doxorubicin, carboplatin, and paclitaxel were used to treat the patient cultures in 3D organoids at indicated doses. In the five ER+ patient cultures (Patient##5, Patient#6, Patient#7, Patient#8, and Patient#9), all three chemotherapies increased the proportion of ALDH+/CD44+ CSL cells versus live cells (CSL cells/Live) (Fig. 1B). This was confirmed with ALDH+/CD44+ CSL cells versus total cells (CSL cells/Total) (Fig. S2B), although the CSL cells versus live cells and versus total cells were reduced by carboplatin in Patient#6. Interestingly, for three TNBC patient samples, the mean values of ALDH, the ALDH+/CD44+ CSL cells versus live cells and total cells were generally decreased. These results suggest that there is heterogeneity in how chemotherapy impacts CSL state. Further, the results from patient tumor cells show that chemotherapy induces CSL traits in ER+ breast cancer.

### Chemotherapy-induced CSL states are recapitulated in breast cancer cell lines

To further investigate these phenomena, we tested four ER+ breast cancer cell lines (CAMA-1, T47D, MCF-7, ZR-75-1) and five TNBC cell lines (MDA-MB-231, BT549, HCC38, HCC1395, HCC1143) using the same experimental settings. As shown in Fig. 1C and Fig. S2C, chemotherapy increased the abundance of ALDH, CD44, CSL cells/Live, and CSL cells/Total in most ER+ cell lines, although T47D and MCF7 have been reported as ALDH-negative [25]. This promotion was less present in TNBC cell lines following chemotherapy, and showed only increased abundance of CD44. The CSL enrichment by chemotherapy in ER+ cells and TNBC is shown by relative mean values of ALDH and CD44, which are highlighted for ER+ patient cells (Fig. S3A), TNBC patient cells (Fig. S3B), ER+ cell lines (Fig. S3C), and TNBC cell lines (Fig. S3D). Spots for TNBC patients and cell lines showed a lesser extent of primitive cell traits compared to ER+ cells. These studies indicate that chemotherapy can lead to enrichment in CSL cells and ER+ cells in particular.

### CSL phenotypes are reversed with an HDAC inhibitor

We sought to find an inhibitor that could reverse chemotherapy induced CSL cells. We used an ER+ breast cancer cell line (CAMA-1; ER+, PR+, and HER2-), a HER2+ and ER-breast cancer cell line (SKBR3; ER-, PR-, and HER2+), and two TNBC cell lines (MDA-MB-231 and BT549) to test for CSL reversal using a 3D organoid culture model for the chemotherapy induced CSL state. The potential CSL inhibitors were selected based multiple lines of data: 1) potential strategies to suppress CSCs summarized by Lin et al.[14] and others, which include targeting pathways that regulate EMT such as hedgehog (vismodegib), Wnt/β-catenin (ICG-001) or CSC markers (disulfiram as an ALDH inhibitor), 2) targeting histone deacetylases which may modulate stem-like signaling pathways (belinostat and entinostat), 3) growth factor receptor pathway inhibitors such as AXL inhibitors (BGB324 and TP-0903), FGFR inhibitor (AZD4547), EGFR inhibitors (afatinib and gefitinib), and 4) cyclin-dependent kinases (CDKs) inhibitors (senexin A, ribociclib and abemaciclib), which are reported to play indispensable roles in processes of stem cell self-renewal [26].

For CSL inhibitor selection, 3D organoids of each breast cancer cell line were treated with chemotherapy for 72 hours, followed by belinostat for another 72 hours, and subjected to ALDEFLUOR/CD44/DAPI staining for FACS analysis. As shown in Fig. 2A, B and Fig. S4A, B, the pan-HDAC inhibitor belinostat consistently reversed both ALDH and CD44 levels promoted by three chemotherapy treatments in all cell lines, while another HDAC inhibitor entinostat, which specifically targets to Class I HDAC (1 and 3), only exhibited reversal effect in HER2+ breast cancer cell line SKBR3. The AXL inhibitor TP-0903 showed some reversal potential in a subset of the cell lines, but not a consistent reversal. Additionally, an increased cytotoxicity of belinostat and TP-0903 were also observed in the viability heat maps for both cell lines at the doses for CSL reversal effect (Fig. 2A, B and Fig. S4A, B). Taken together, these data indicate that belinostat may serve as a general anti-chemo-induced CSL modifier in breast cancer.

**Fig. 2.**
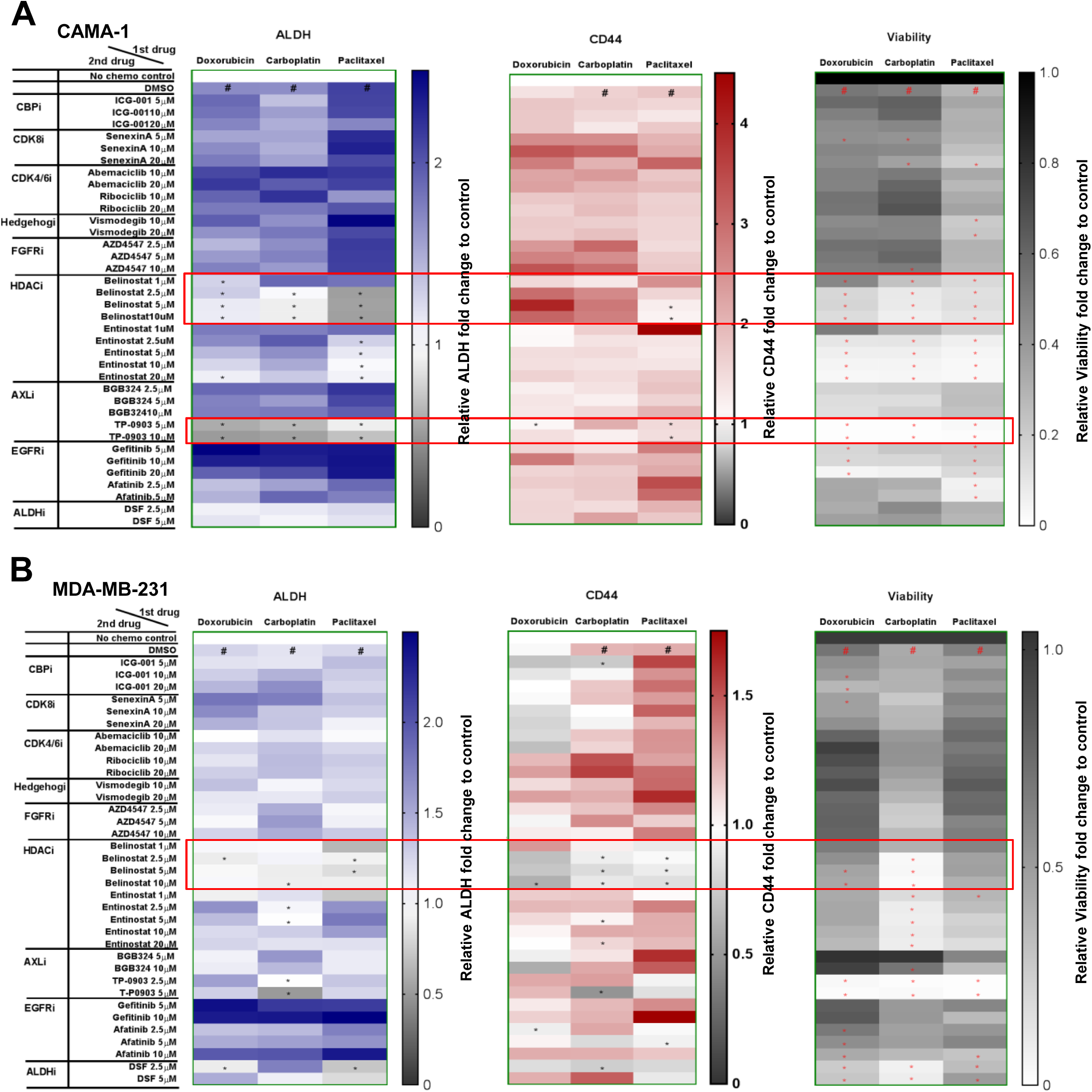
Belinostat reduced chemotherapy-promoted CSL state. Heat maps of FACS mean values for CSL reversal drugs screening with CAMA-1 (**A**) and MDA-MB-231 (**B**) cell lines. Cells were cultured in medium (3D) and treated for 72 hours with doxorubicin (0.1µM for CAMA-1, 0.5µM for MDA-MB-231), carboplatin (50µM for all cell lines), and paclitaxel (1µM for all cell lines), and then treated with potential CSL inhibitors as indicated doses for 72 hours before ALDEFLUOR/CD44 staining. The FACS mean values of ALDH and CD44 in live populations and viability were shown in the heat maps. The vehicle controls (DMSO+DMSO) were set as fold one. All the other chemotherapy plus DMSO/inhibitor combinations were expressed as the fold change relative to the controls. The Chemo plus DMSO served as the positive controls. The mean values of triplicate tests were showed with color indicated as the legends. In heat maps of ALDH and CD44, # is chemo plus DMSO significantly higher (p<0.05) than no treatment control, and in heat maps of viability, # is chemo plus DMSO significantly lower (p<0.05) than no treatment control. In all heat maps, * is chemo plus inhibitor significantly lower (p<0.05) than chemo plus DMSO. The potential inhibitors are highlighted with red blocks as its reversal effect for both ALDH and CD44.

### Belinostat, a pan-HDAC inhibitor, reverses chemotherapy-derived CSL enrichment in ER+ breast cancer cells

To further evaluate the potential of belinostat to enact chemo-induced CSL reversal, we performed a time-course FACS analysis from Day 1 to Day 6 to measure the CSL phenotype every day using the experiment strategy shown in Fig. 3A. For the ER+ breast cancer cell line CAMA-1, as shown in Fig. S5A, FACS analysis showed steadily increasing ALDH levels with all three chemotherapies (doxorubicin, carboplatin, and paclitaxel) from Day 1 to Day 6, while the CD44 levels showed steady increases from Day 1 to Day 6 with doxorubicin and paclitaxel. Overall, there was an increase in the CSL cells versus live cells from Day 1 to Day 6 in all three chemotherapy groups (Fig. 3B). The overall viability continuously decreased each day following all three chemotherapy treatments, while belinostat further reduced viability compared to control from Day 4 to Day 6 (Fig. 3B), indicating an increase in cytotoxicity with belinostat treatment in addition to its CSL reversal effect. Importantly, belinostat reversed the upward trajectory of ALDH (Fig. S5A), CSL cells versus live cells (Fig. 3B), and CSL cells versus total cells (Fig. S5A) with a significant reduction in treated versus control at Day 5 and Day 6. Of note, CD44 was only reversed by belinostat following paclitaxel treatments, but not carboplatin and doxorubicin treatment (Fig. S5A), suggesting some differences in chemotherapy effects. Summarized results across chemotherapy treatments show a statistically significant reduction in ALDH, CSL cells versus live cells, and CSL cells versus total cells with belinostat treatment at Day 6 (Fig. 3B and Fig. 5A).

**Fig. 3.**
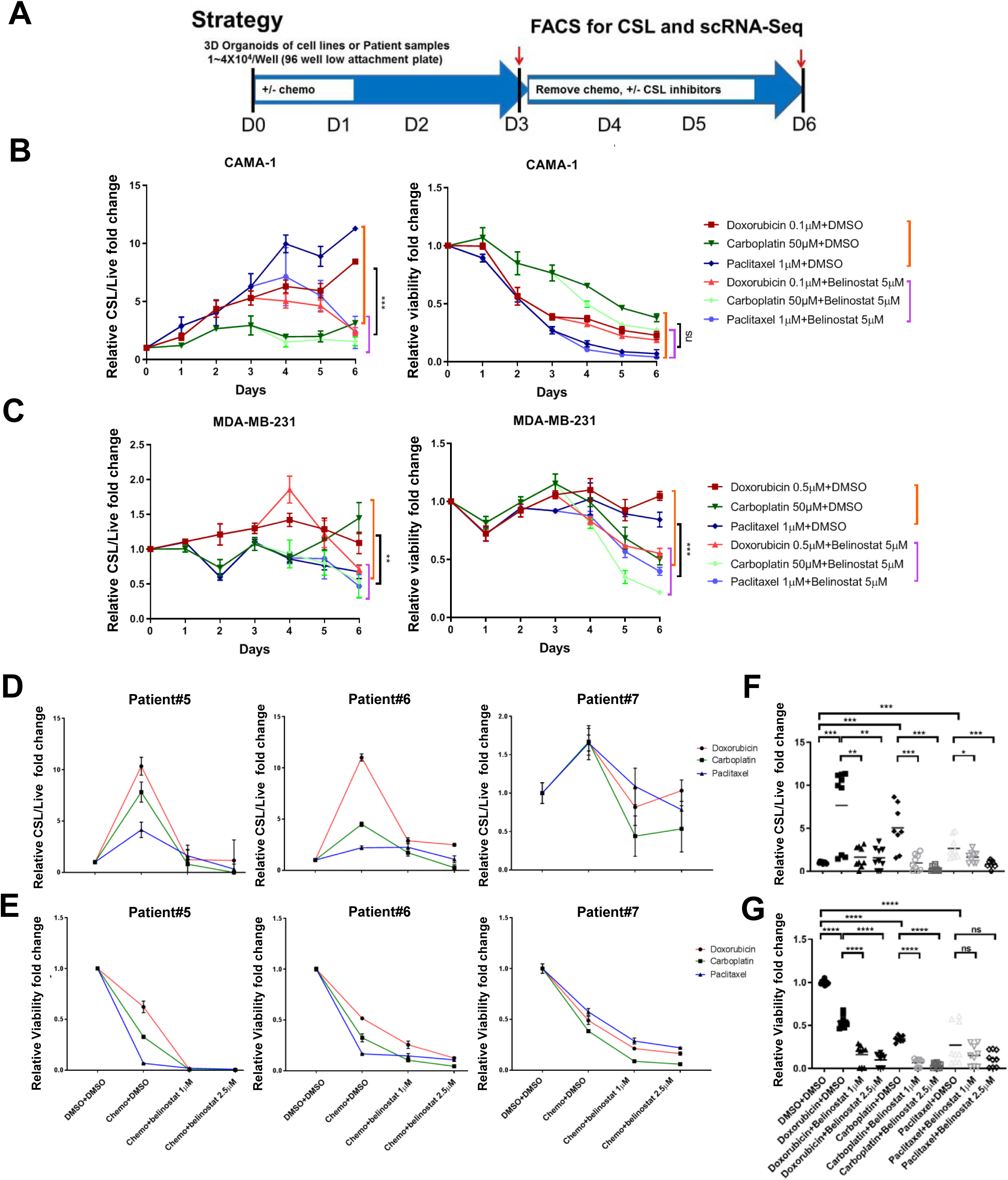
Belinostat inhibited chemotherapy-promoted CSL state in ER+ cell lines and patient cancer cell samples. (**A**) Cell culture strategy for CSL reversal drug screening with cancer breast cell lines after chemotherapy promotion. The CSL cells/live and viability were shown in (**B**) for CAMA-1 and (**C**) for MDA-MB-231 with curves. CAMA-1 cells and MDA-MB-231 cells were cultured in medium (3D) and treated for 72 hours with doxorubicin (0.1µM for CAMA-1, 0.5µM for MDA-MB-231), carboplatin (50µM), and paclitaxel (1µM), and then replaced with medium containing belinostat (5µM) for 72 hours incubation. Samples were collected every day from Day 1 to Day 6, and followed by ALDEFLUOR/CD44/DAPI staining and FACS analysis. The differences between chemo plus DMSO (9 replicates from three chemo plus DSMO) and chemo plus belinostat (9 replicates from three chemo plus belinostat) at Day 6 were evaluated by student’s T tests. (**D**) The CSL cells/live and (**E**) viability were expressed with curves for Patient#5, #6, and #7. Patient cells were cultured in Renaissance medium (3D) and treated with chemotherapy (doxorubicin 0.1 µM, carboplatin 0.1mM, paclitaxel 1µM) for 72 hours, and then treated with DMSO/belinostat (1µM or 2.5 µM) for another 72 hours, followed by ALDEFLUOR/CD44/DAPI staining and FACS analysis. Column graphs with individual values were generated by combining 9 replicates from three patients for each treatment to evaluate the changes in CSL cells/live (**F**) and viability (**G**). In (**B**-**G**), DMSO treatment only in each day served as controls and was set as fold one. All the other treatments were expressed as the fold values relative to the controls. In (**B**-**E**), each spot represents the mean values of triplicate tests. The statistical analysis was performed with student’s t-tests between the indicated groups. Significance is marked with * for P<0.05, ** for P<0.01, *** for P<0.001, **** for P<0.0001.

**Fig. 4.**
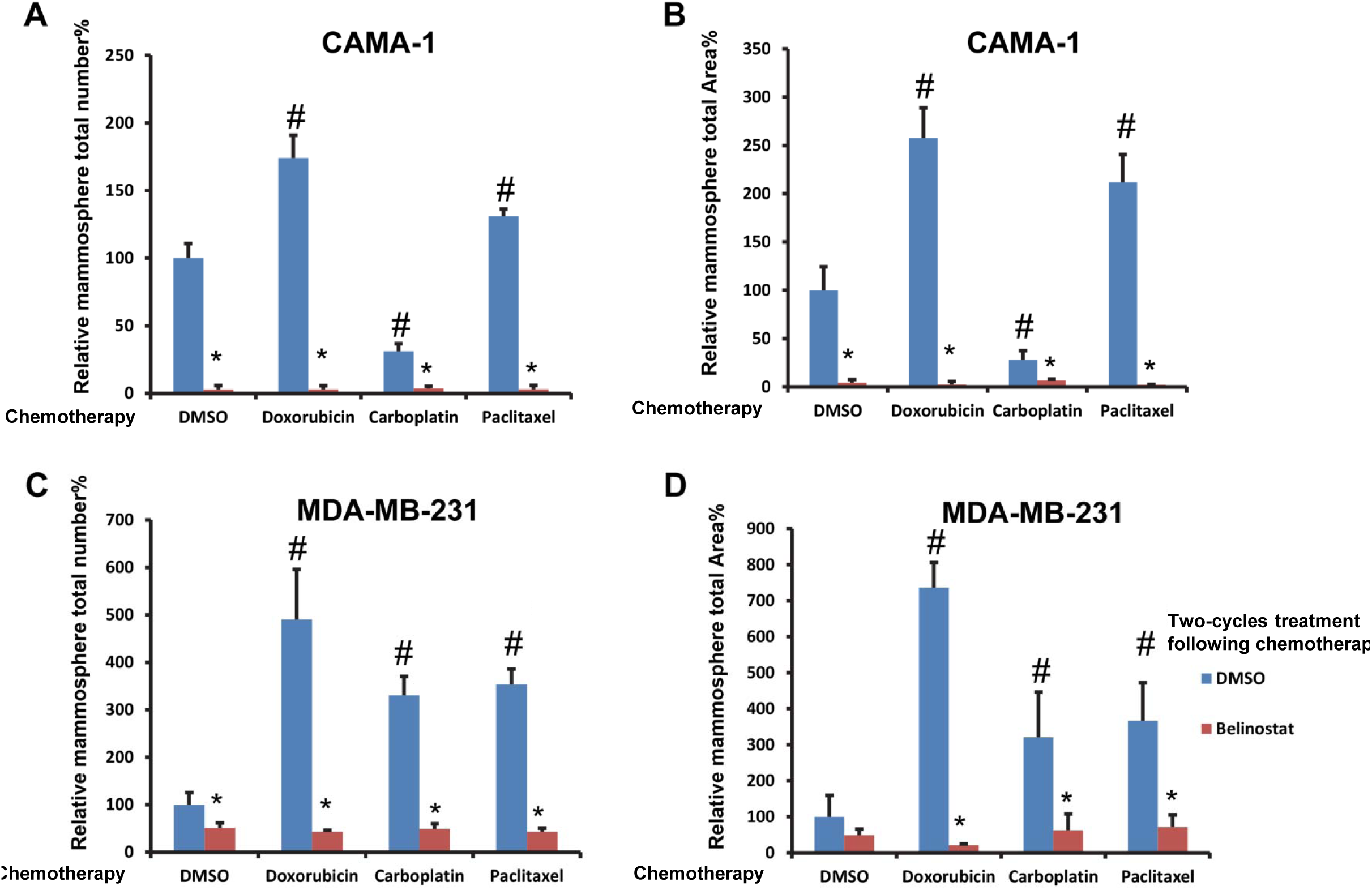
Belinostat inhibited mammosphere formation of chemotherapy-treated breast cancer cell lines. CAMA-1 and MDA-MB-231 cells from spheroids with 3 days chemo treatment (DMSO, doxorubicin 0.1µM for CAMA-1, 0.5µM for MDA-MB-231, carboplatin 50µM, and paclitaxel 1µM) were collected, dissociated, and cultured for two 14 days periods with mammosphere assay medium containing DMSO/belinostat (5µM) as described in the Methods. Mammosphere total numbers and total area of CAMA-1 are shown in (**A**) and (**B**). Mammosphere total numbers and total area of MDA-MB-231 are shown in (**C**) and (**D**). The treatments with DMSO only serve as the control in each figure and set as fold one. All the other treatments are expressed relative to the controls in the figures. The statistical analysis was performed with student’s t-tests between the indicated groups with triplicate tests. # is p<0.05 relative to DMSO plus DMSO only, * is p<0.05 relative to chemo plus DMSO.

**Fig. 5.**
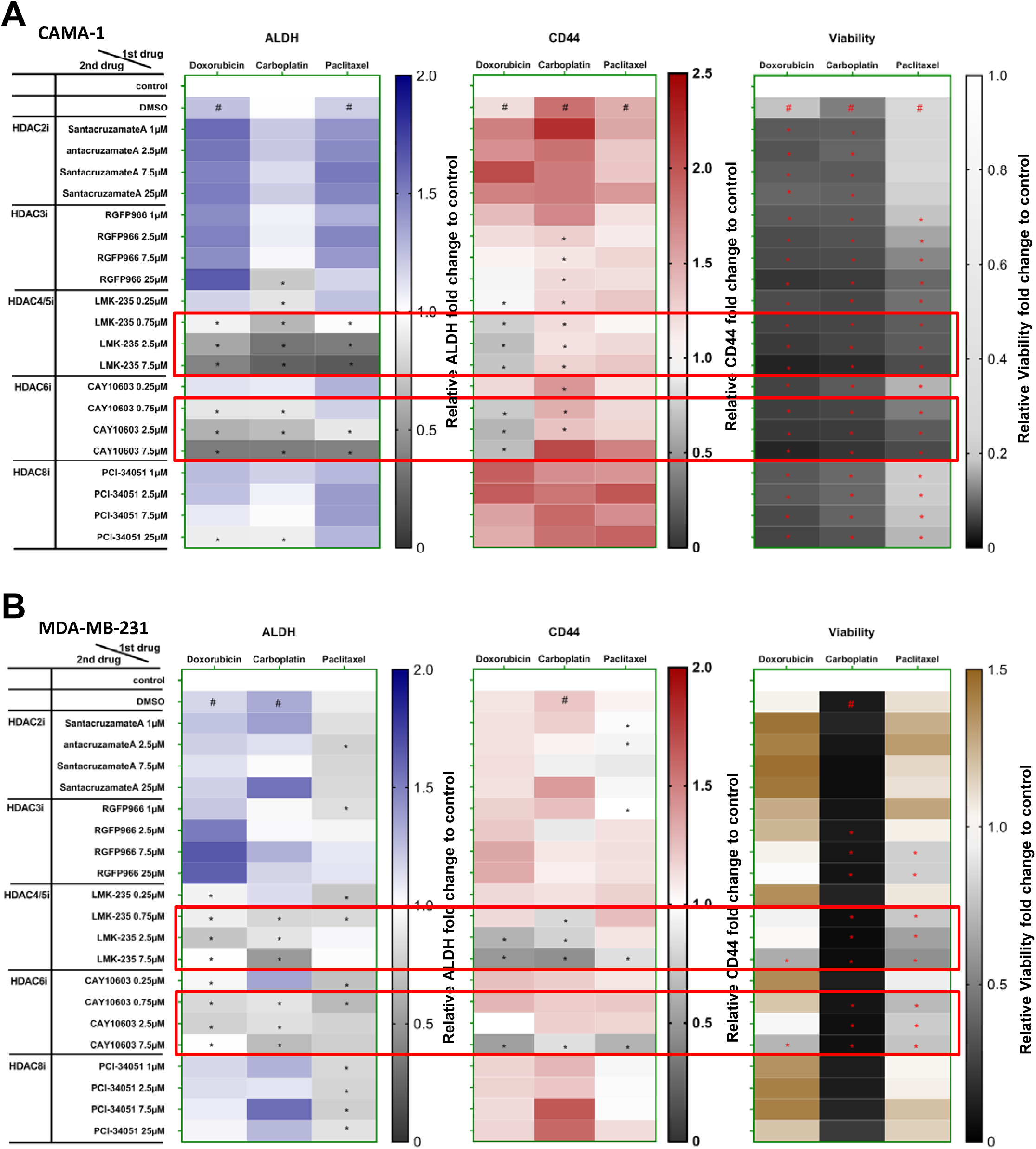
Specific HDAC isoforms target chemotherapy-acquired CSL state. Heat maps of FACS mean values for CSL reversal drugs screening with CAMA-1 (**A**) and MDA-MB-231 (**B**) cell lines. Cells were cultured in medium (3D) and treated for 72 hours with doxorubicin (0.1µM for CAMA-1, 0.5µM for MDA-MB-231), carboplatin (50µM for all cell lines), and paclitaxel (1µM for all cell lines), and then treated with specific HDAC isoforms inhibitors as indicated doses for 72 hours before ALDEFLUOR/CD44 staining. The FACS mean values of ALDH and CD44 in live populations and viability were shown in the heat maps. The vehicle controls (DMSO+DMSO) were set as fold one. All the other chemotherapy plus DMSO/inhibitor combinations were expressed as the fold change relative to the controls. The Chemo plus DMSO served as the positive controls. The mean values of triplicate tests were showed with color indicated as the legends. In heat maps of ALDH and CD44, # is chemo plus DMSO significantly higher (p<0.05) than no treatment control, and in heat maps of viability, # is chemo plus DMSO significantly lower (p<0.05) than no treatment control. In all heat maps, * is chemo plus inhibitor significantly lower (p<0.05) than chemo plus DMSO. The potential specific HDAC isoforms inhibitors target to CSL cells are highlighted with red blocks as its reversal effect for both ALDH and CD44.

Less consistent results were found in MDA-MB-231, a TNBC cell line (Fig. 3C and Fig. S5B). CD44 (Fig. S5B), and CSL cells versus live cells (Fig. 3C) slightly increased with doxorubicin and carboplatin, but not paclitaxel, while ALDH and CSL cells versus total cells were not generally increased by chemotherapy treatments. Further, ALDH, CD44, CSL cells versus live cells, and CSL cells versus total cells (Fig. 3C and Fig. S5B) all decreased with belinostat following all three chemotherapy treatments. Belinostat treatment exhibited a statistically significant reduction across all chemotherapies in ALDH, CD44, CSL cells versus live cells, CSL cells versus total cells, and viability at Day 6 (Fig. 3C and Fig. S5B). Overall, the CSL reversal following chemotherapy was less pronounced in TNBC than ER+ cells, which was indicated by the reduction of CSL cells versus live cells in MDA-MB-231 (0.67-1.4 folds in chemotherapy plus DMSO drop to 0.46-0.71 folds in chemotherapy plus belinostat) versus those in CAMA-1 (3.1-11.2 folds in chemotherapy plus DMSO drop to 1.56-2.4 folds in chemotherapy plus belinostat), perhaps reflecting the lack of chemotherapy-promoted CSL traits in TNBC versus ER+ cells, and higher baseline CSL in TNBC [27].

As chemotherapy strongly promoted CSL state in ER+ breast cancer patient samples, with a less pronounced effect in TBNC patient samples as indicated above (Fig. 1B), we further tested the CSL reversal effect of belinostat in ER+ patient cancer cells. These studies use patient samples to further test the cell line results from above. Three ER+, PR+, Her2-breast cancer patient samples (Patient#5, Patient#6, and Patient#7) were used to perform the same experiments as used for the cell lines. The patient 3D organoids were treated with doxorubicin, carboplatin and paclitaxel for 72 hours respectively, and followed by either control DMSO or belinostat (1µM or 2.5µM) for 72 hours. Relative fold changes compared to DMSO control were expressed with curves for each individual patient sample treated with three chemotherapies. As shown in Fig. S6A and Fig. 3D, FACS analysis of these cells showed that ALDH and CSL cells versus live cells were generally increased by chemotherapy and reversed by belinostat. Note, CD44 decrease was found in the majority but not all of the patient cells (Fig. S6B). CSL cells versus total cells were reduced by belinostat in the patient samples, although induction of CSL state alter chemotherapy varied by patient, most likely due to prior treatments the patient received and various levels of response to chemotherapy (Fig. S6C). Importantly, we found all chemotherapies decreased the viability of cancer cell, and belinostat further decreased viability in a dose-dependent manner (Fig.3E). Across all patient samples, FACS analysis showed increases in ALDH (Fig. S6D), CSL cells versus live cells (Fig. 3F), and CSL cells versus total cells (Fig. S6E) following chemotherapy treatment compared to vehicle controls, and all are significantly decreased by belinostat in dose-dependent manner compared to chemotherapy treatment only controls. Note, CD44 levels did not show a significant common effect either on chemotherapy promotion or belinostat reversal, although the expected trends could be observed between indicated groups (Fig. S6F). The viability decreased with all chemotherapies, and further decreased with belinostat in dose-dependent manner (Fig. 3G). Overall, these experiments validated the cell line studies with patient tumor samples and show that belinostat may serve as a general anti-CSL inhibitor following multiple chemotherapies in ER+ breast cancer, reflecting a one-two punch treatment strategy.

### Belinostat inhibits chemotherapy induced mammosphere formation

We next tested the ability of belinostat to modulate CSL self-renewal and proliferation using a mammosphere formation assay [28]. Two cycles of mammosphere culture were performed to select stem-like cells with self-renewal ability and eliminate various transitions following chemotherapy +/- belinostat. Both ER+ (CAMA-1) cells and TNBC (MDA-MB-231) cells how a significantly diminished mammosphere formation following belinostat treatment after chemotherapy after two-cycle cultures (Fig. 4A, B, C, D). The majorities of chemotherapies increased the total mammosphere numbers and areas compared to DMSO control only, while belinostat decreased the total mammosphere counts as well as their areas compared to the chemotherapy plus DMSO control.

For the ER+ cells, doxorubicin and paclitaxel both increased mammosphere counts (up to 174.1% and 131.1%), and mammosphere total areas (up to 257.9% and 211.9%) compared to DMSO control (set to 100% for each condition). Strikingly, belinostat dramatically reduced the mammosphere counts (to 2.9-3.7%) and mammosphere total areas (to 2.3-6.7%) following all chemotherapy treatments. Similarly, for the TNBC cells, all chemotherapies strongly increased mammosphere counts (up to 490.6%, 330.8%, and 353.8%), and mammosphere total areas (up to 736.2%, 320.8%, and 366.6%) compared to DMSO control. Belinostat again reduced the mammosphere counts (42.7-51.3%) and mammosphere total areas (21.5-71.9%) following all chemotherapy treatments. Thus, the general anti-CSL effect of belinostat is supported by its ability to inhibit CSL self-renew and proliferation with mammosphere assays.

### HDAC class II regulate chemo-acquired CSL reversal

As the pan-HDAC inhibitor belinostat, but not the Class I HDACs inhibitor, entinostat [29,30] reversed CSL states following chemotherapy treatments, specific HDAC targets were tested in order to define HDAC proteins essential for this activity. HDAC inhibitors targeting specific HDAC isoforms were used to test for CSL reversal activity using the FACS strategy described above (Fig. 2). Tested were Class I HDACs inhibitors SantacruzanateA (HDAC2i) [31], RGFP966 (HDAC3i) [30], PCI-34051 (HDAC8i) [29]; Class IIa HDACs inhibitor LMK-235 (HDAC4/5i) [32]; and Class IIb HDACs inhibitor CAY10603 (HDAC6i) [33, 34]. As shown in Fig. 5A, B, the HDAC4/5 inhibitor LMK-235 and HDAC6 inhibitor CAY10603 reversed both ALDH and CD44 levels promoted by three chemotherapy treatments in both cell lines, while other specific HDAC inhibitors did not reverse these primitive traits. These results suggest that HDAC class IIa and IIb (HDAC4/5 and HDAC6), but not HDAC class I, are essential in HDAC mediated CSL traits plasticity.

To further test the CSL state reversal effect of Class II HDACs inhibitors, a time-course FACS analysis was performed with ER+ cell line CAMA-1 (Fig. S7A, B) and TNBC cell line MDA-MB-231 (Fig. S8A, B) treated with LMK-235 or CAY10603 following three chemotherapies using the strategy described in Fig. 3B and C. Both Class II HDAC inhibitors decreased cell viability compared to chemotherapy treatment alone, and exhibited strong CSL reversal activity similar to belinostat, including decreases in ALDH levels and statistically significant diminished levels of both live and total CSL cells following chemotherapy (Fig. S7A, B and Fig. S8A, B). Detailed FACS plots for CSL cell measurements (ALDH vs. CD44) at Day 6 are shown in Fig. S9A and Fig. S9B. These data detail the significant reduction in the number of CSL cells constrained in the ALDH+, CD44+ positive quadrants with chemotherapy treatment followed by class II HDAC inhibitors compared to chemotherapy alone. Interestingly, the treatments show different live CSL cell reversal trajectories, with LMK-235 and CAY10603 exhibiting relatively strong effects on primitive trait reversal. Taken together, these results indicate that class II HDACs (HDAC4/5/6) regulate chemo-acquired stemness states.

### MYC pathways mediated CSL promotion/reversal upon chemotherapy/belinostat treatment

To understand how belinostat reverses the chemotherapy-induced CSL properties, we performed single-cell RNA-Seq analysis with two ER+ patient samples (Patient#5 and Patient#6). We performed an unbiased clustering of the gene expression profiles of the Patient#6 and Patient#5 cells and saw that there are changes in the gene expression profiles of the cells after chemotherapy treatments and combination chemotherapies plus belinostat (Fig. 6A, B). The Patient#6 cells exhibited relatively similar transcriptional programs to all chemotherapies, and also to the combination treatments. Chemotherapy generated a distinctive cell population that was further changed by chemo+belinostat treatment (Fig. 6A). This indicates that, while we previously saw that belinostat can reverse the stem cell phenotypes, it does not reverse all aspects of the chemotherapy. In contrast, the Patient#5 cells showed a drug-specific response where each chemotherapy led to distinct changes in the cell populations, and relatively little changes in the transcriptional programs were induced by belinostat (Fig. 6B).

**Fig. 6.**
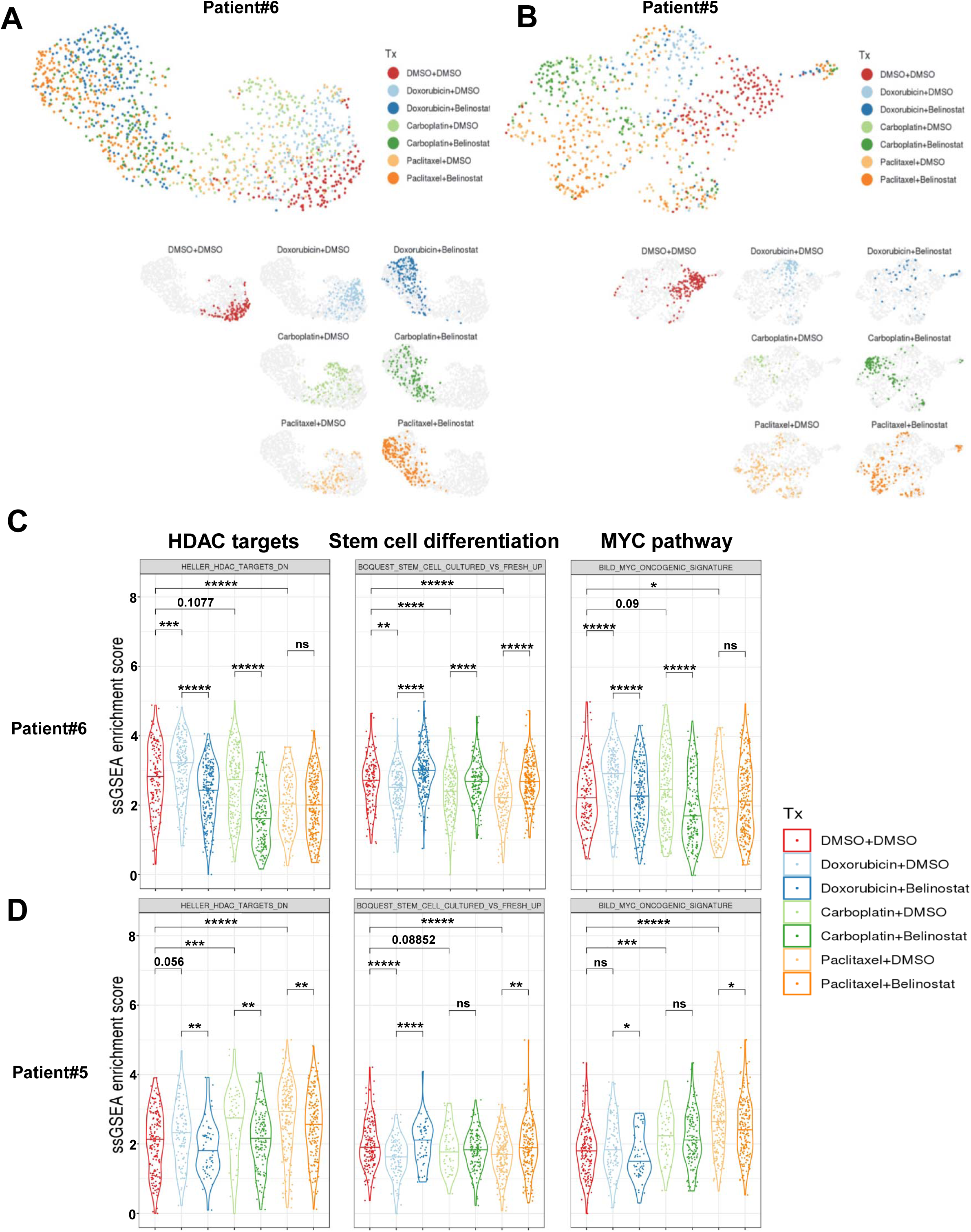
Chemotherapy and belinostat reprogrammed transcriptome of patient cells. Unbiased UMAPs for ER+ patient sample Patient#6 (**A**) and Patient#5 (**B**) under treatment (Tx). Patient cells were cultured in Renaissance medium (3D) and treated with chemotherapy (doxorubicin 0.1 µM, carboplatin 0.1mM, paclitaxel 1µM) for 72 hours, and then treaded with DMSO/belinostat (1µM) for 72 hours. The cells were collected, purified with Dead Cell Removal Kit, and subjected to ICELL8® Single-Cell System as described in Methods. For each patient, the UMAPs for all treatments were shown in the top, and the UMAPs for each individual treatment were shown in the bottom with different colors indicated as the legends. Violin plots with GSEA pathways enrichment scores related to HDAC targets, stem cell differentiation and MYC pathway were presented in (**C**) for Patient#6 and (**D**) for Patient#5. The statistical analysis was performed with student’s t-test between the indicated groups, and significance is marked with * for P<0.05, ** for P<0.01, *** for P<0.001, **** for P<0.0001.

As belinostat can induce changes in the stem cell related markers (Fig. 3), the lack of obvious changes in the overall transcriptional profiles suggest that the differences in the stem cell program may leave a relatively small imprint on the overall profiles or that progenitor cell transcriptional profiles are more nuanced. To focus on the specific (rather than overall) changes seen with drug treatment, we performed a pathway analysis to identify the pathways that are altered by chemotherapy or combination treatment.

Shared reversal pathways were selected as described in Methods, and heat maps were generated based on the averages of ssGSEA enrichment scores for the shared reversal pathways for both Patient#6 (Fig. S10A, B) and Patient#5 (Fig. S10C, D). For each patient sample, we selected the pathways that were either increased by chemo and decreased by belinostat or the inverse pattern (Fig. S10). In our shared reversal pathways analysis, we found that HDAC pathways (HELLER_HDAC_TARGETS_DN) were up-regulated by all three chemotherapy treatments in both patient samples and down-regulated by belinostat except paclitaxel group in Patient#6 (Fig. 6C, D). This indicates that chemotherapy treatments promoted HDAC targets in this gene set (eg, CD44 as stemness marker, BCL2 as anti-apoptotic regulator), while belinostat, a pan-HDAC inhibitor (HDACi), reversed them; concomitant with the changes in CSL properties. This is consistent with prior reports showing that HDAC expression is up-regulated in CSCs [13] and HDAC inhibitors can inhibit CSL phenotypes through its inhibition of acetylation activity [13, 14].

We also observed that a stem cell differentiation signature (BOQUEST_STEM_CELL_CULTURED_VS_FRESH_UP) was reduced by all three chemotherapy treatments and promoted by belinostat in both patient samples (Fig. 6C, D). In another words, the CSL state of the survival single cells were up-regulated by all chemotherapy treatments, and down-regulated by belinostat. Consistent with CSL state, the MYC pathway (BILD_MYC_ONCOGENIC_SIGNATURE) was increased by most of chemotherapy treatments, and inhibited by belinostat except paclitaxel group in both Patient#6 (Fig. 6C) and Patient#5 (Fig. 6D). Belinostat has been reported to regulate MYC expression transcriptionally and post-transcriptionally through regulation of MYC acetylation [35]. MYC has previously been shown to be a factor in CSC reprogramming and self-renewal in basal-like breast cancer [36], and may contribute to reversal of the CSL promotion-reversal process mediated by chemotherapy and HDACi.

## Discussion

Cancer stem cells (CSCs) have the capacity to initiate growth of new tumors as well as cause disease relapse; further, these cells are not targeted by conventional chemotherapy or other treatment strategies [37]. Therefore, there remains potential in developing treatment approaches that minimize CSC or CSL cells to delay or prevent refractory cancer development. Our study indicates that subclonal selection in patients during therapy promotes CSL state in breast cancer. Further, both flow cytometry and mammosphere assays show a reversal of CSL state following chemotherapy by the use of the HDAC inhibitor belinostat. Single-cell RNA sequencing analysis indicated that the RNA transcriptome was strongly reprogramed by chemotherapy treatments, and reversed by belinostat, which promoted stem cell differentiation, and inhibited HDAC targets and MYC pathway. The imbalance between acetylation and deacetylation of histone or non-histone proteins, which are regulated by histone acetyltransferases (HATs) and deacetylases (HDACs), have been proved closely to be associated with cancer development [38]. Acetylation and deacetylation of proteins is involved in various stemness-related signal pathways and can regulate self-renewal and differentiation signaling [38]. The pan HDAC inhibitor used in this study has been used previously for their anti-tumor and anti-CSC activity by targeting multiple signaling pathways at different molecular levels [14, 39]. Besides chromatin remodeling via histone acetylation, HDAC inhibitors could also block key signaling pathways pertinent to CSC maintenance, including HIF-1α, Stat3, Notch1, β-catenin, NF-κB, MYC, and c-Jun with the ability of different HDAC isoforms to regulate protein stability and/or activity [14].

Although HDAC inhibitors have been used alone or in combination therapy to suppress the CSC population in different settings [14], how HDAC inhibitors reverse CSL state following standard of care chemotherapies is not well-defined. Previous clinical trials of HDACi target to CSL cells either focused on the monotherapy for refractory peripheral T-cell lymphoma, or concurrent combination of chemotherapy with HDACi on solid tumors [39, 40]. However, these clinical trials and research assessed antitumor activity by apoptosis/viability test, more than targeting to acquired stemness. In addition, it has not been studied if specific chemotherapies or this class of drug generally, can induce CSL state. Further, it has not been studied if HDAC inhibitors can reverse CSL states driven by different chemotherapies. As detailed results above, our study addresses subclonal evolution of stem-like traits in patients as they acquire resistance to chemotherapy, as well as approaches to reverse this acquired resistant trait.

In this study, we initially tested multiple drugs published previously to block stem cell-like state. Only one was effective following chemotherapy induced stem-like states across cell lines and patient tumor cells. Although there was some heterogeneity in ALDH or CD44 expression level modulated by different chemotherapies, we generally saw an increase in CSL levels for ER+ tumor cells and to a lesser extent TNBC cells. Belinostat broadly blocked increased CSL cells following chemotherapy treatment based on flow cytometry and mammosphere assays. Although HDAC inhibitors have been proved for their epigenetic effect on the reprogramming of gene expression in cancer cells, which leads to growth arrest, differentiation, and apoptosis [41], their roles in suppressing the self-renewal capability and driving the differentiation of CSCs could also enhance the cancer cell sensitivity to chemotherapy [38], which might lead a synergistic effect for CSL elimination. HDAC inhibitors also prevent or reverse the development of drug resistance by its inhibition on drug transporters (MDR1) expression, thus augmenting the effect of combined therapy with their synergism [42]. The mammosphere formation assays in our study demonstrated that belinostat could abolish the self-renewal capability of survival cells after chemotherapy treatments, which has resulted in resistant CSL subtypes. Importantly, these results were consistent across different chemotherapies tested, suggesting a convergence on CSL resistance. The synergistic effects of these drug combinations as well as optimal timing and sequence should be determined by further experiments and mathematical modeling.

In our FACS strategy to identify the CSL cells, ALDH and CD44 were selected as the CSL state markers, rather than CD24. Of Note, the initial state of CAMA-1 starts from very low primitive cell traits (indicated by low CD44 level) compared to MDA-MB-231, which might suggests more potential for CSL state induction for ER+ cells other than TNBC. Recently, two types of distinct CSCs, epithelial-like (E) ALDH+ CSCs and mesenchymal-like (M) CD44+/CD24− CSCs, have been identified [37]. Liu et al reported that E and M CSCs co-existed in all breast cancer subtypes, but their proportions were varied [43]. ALDH+ CSCs located within the core of the tumor with high expression of E-cadherin but low expression of vimentin and ZEB1, undergo proliferation at hypoxia conduction, while CD44+/CD24− CSCs located at the edge of the breast tumor with low expression of E-cadherin but high expression of vimentin and ZEB1, undergo migration and invasion [43]. These two subtypes of CSCs exhibit differential signal pathway regulations, plasticity, and differently responses to treatment. E and M CSCs were also found to be interconvertible in EMT-MET signaling. Therefore, co-inhibition of both E and M subtypes of CSCs may be needed to develop effective therapy [43]. The detailed FACS analysis following chemotherapies (Fig. S9A, B) suggests belinostat potentially exhibited dual inhibition of E and M CSL cells, with its reversal effect on ALDH and CD44 in TNBC cell line MDA-MB-231 (initially ALDH low/CD44 high), which is enriched with M subtype of CSL cells [27], and on ALDH in ER+ cell line CAMA-1 (initially ALDH high/CD44 low) and patient samples, which is enriched with E subtype of CSL cells. Compared to the FACS analysis for CSL state, the mammosphere assay tests the self-renewal ability of resistant cells rather than the changes of stemness makers. Cells with high level expression of CD44 are more resistant to chemotherapy and likely to form mammosphere [27], which would benefit the TNBC than ER+ cells due to their enrichment of progenitor cells (ALDH negative, CD44 high). In this case, inhibition effect of belinostat on CD44+ cells could play a major role to abolish the self-renewal capability of chemo-resistant cells. Reversal of CD44 in ER+ cell lines/patient samples and inhibition of ALDH in TNBC cell line were also identified by FACS analysis, which varied among cell lines/patient samples with different chemotherapies. Thus, future research may help delineate cell populations sensitive to belinostat treatment.

In the single-cell RNA sequencing analysis, we are interested in visualizing the pattern of activation of the pathways across the different treatment groups, other than seeking the overall changes by leveraging multiple pairwise comparisons against control treated cells and then plotting the enrichment score or a metric based on the statistical significance. The approach we used could more directly provide information such as the relative magnitude of the changes, as well as the baseline levels of activation in the control group. In the single-cell RNA sequencing analysis presented here, chemotherapy followed by belinostat revealed that MYC could be an important mediator for these two distinct subtypes of CSCs. MYC induces the dedifferentiation towards a progenitor-like state by epigenetic reprogramming [44]. Further, mammosphere with MYC overexpression showed enrichment for cells expressing ALDH1 [45]. MYC-dependent metabolic reprogramming is related to CD44 variant-dependent redox stress regulation in CSCs [44]. Targeting MYC family by HDAC inhibition, such as valproic acid, could contribute to the down-regulation of MYC, in turn to up-regulate of CDKN1A/B (p21/CIP1/WAF1, p27/KIP1) to induce cell cycle arrest and autophagic cell death [44, 46].

HDACs can be classified into four classes (class I to IV): Class I (HDAC1, HDAC2, HDAC3, and HDAC8). Class IIa (HDAC4, HDAC5, HDAC7, HDAC9) and IIb (HDAC 6 and 10), class III (sirtuins 1-7), and class IV (HDAC11) [38]. Class I HDACs only localize in nucleus, while Class II HDACs exhibit their ability to shuttle between nucleus and cytoplasm, and can deacetylate non-histone proteins in cytoplasm [38, 47]. Positive associations of high expression of ClassII HDACs with cell proliferation and CSC differentiation were found in various cancer types, including gastric, osteosarcoma, and colon cancer with HDAC4 [48, 49], breast and lung cancer with HDAC 5 [50, 51], and anaplastic thyroid, breast cancer and glioblastoma multiforme (GBM) with HDAC6 [52, 53, 54]. HDAC5 and HDAC6 transcriptional regulation on MYC has been reported to play a key role in these processes [34, 52, 54]. The HDAC4/5i LMK-235 and HDAC6i CAY10603 both exhibited their CSL reversal effect on reducing CSL abundance and additional cytotoxicity, suggesting their strong potential on clinical therapy to eliminate the chemo-promoted CSL cells. As the pan-HDAC inhibitors have been reported for their side effects [55], the Class II HDAC inhibitors targeting to primitive traits, might become better choices in the combination clinical therapy in the future.

The patient data described in this study supports the model that tumor subclones with enrichment of a CSL phenotype are selected during acquisition of a drug resistant state. It remains unclear how the genetic composition of tumor subclones impacts phenotype. The CNVs used to identify these survivor subclones harbor numerous genes, which may combine to enhance plasticity or a CSL state. This question could be further elucidated in future studies with a larger sample size.

## Conclusion

Taken together, our findings demonstrated inhibition on HDACs, especially Class II HDACs (HDAC4/5/6), reverses chemotherapy-induced acquisition of primitive cell traits and drug-resistance in ER+ breast cancer, and provided an effective ‘one-two punch’ strategy to combat refractory ER+ breast cancer with sequential chemotherapy and HDACi combination.

## Abbreviations

ER: estrogen receptor;
PR: progesterone receptor;
HER2: human epidermal growth factor receptor 2;
TNBC: triple negative breast cancer
CSC: cancer stem cell
CSL: cancer stem cell-like;
ALDH: Aldehyde dehydrogenases
scRNA-Seq: single cell RNA sequencing
HDAC: Histone deacetylase
HDACi: Histone deacetylase inhibitor
ssGSEA: Single-sample Gene Set Enrichment Analysis
CNV: Copy number variations
WGS: whole genome sequencing
WES: whole exome sequencing
EMT: Epithelial to mesenchymal transition
MET: mesenchymal-to-epithelial transition
CDK 4/6: Cyclin-Dependent Kinase 4/6
CDK 8: Cyclin-Dependent Kinase 8
FGFR1: Fibroblast growth factor receptor
EGFR: epidermal growth factor receptor
AXL: Axl receptor tyrosine kinase
CBP: CREB-binding protein

## Declarations

### Ethics approval and consent to participate

Informed consent was obtained from all patients in this study. Protocols were approved by the University of Utah Institutional Review Board and the City of Hope Institutional Review Board.

### Consent for publication

Not applicable

### Availability of data and materials

The scRNA-Seq datasets generated and/or analysed during the current study are available in the Gene Expression Omnibus (GEO) database under accession GSE147326.

### Authors’ contributions

F.C. and S.W.B. contributed study design and coordination, wrote manuscript, data processing, and analysis. J.L., A.N. and J.T.C. contributed single cell-RNA sequencing data processing and analysis. P.A.C. contributed single-cell RNA sequencing preparation. M.S. and P.J.M. coordinated sample collection. J.A.M. contributed to data analysis. J.T.C. and M.K. contributed study design. A.H.B. conceived the study, contributed study design and coordination, wrote manuscript, and performed data analysis. The authors read and approved the final manuscript.

## Acknowledgements

We thank the anonymous patients used in our study for their generous contributions to research. The AXL inhibitor BGB324 was provided by BerGenBio. Research reported in this publication utilized the High-Throughput Genomics and Bioinformatic Analysis Shared Resource at Huntsman Cancer Institute at the University of Utah

