## Supplementary Figures for "A ‘one-two punch’ therapy strategy to target chemoresistance in estrogen receptor positive breast cancer"

**Supplementary Data**

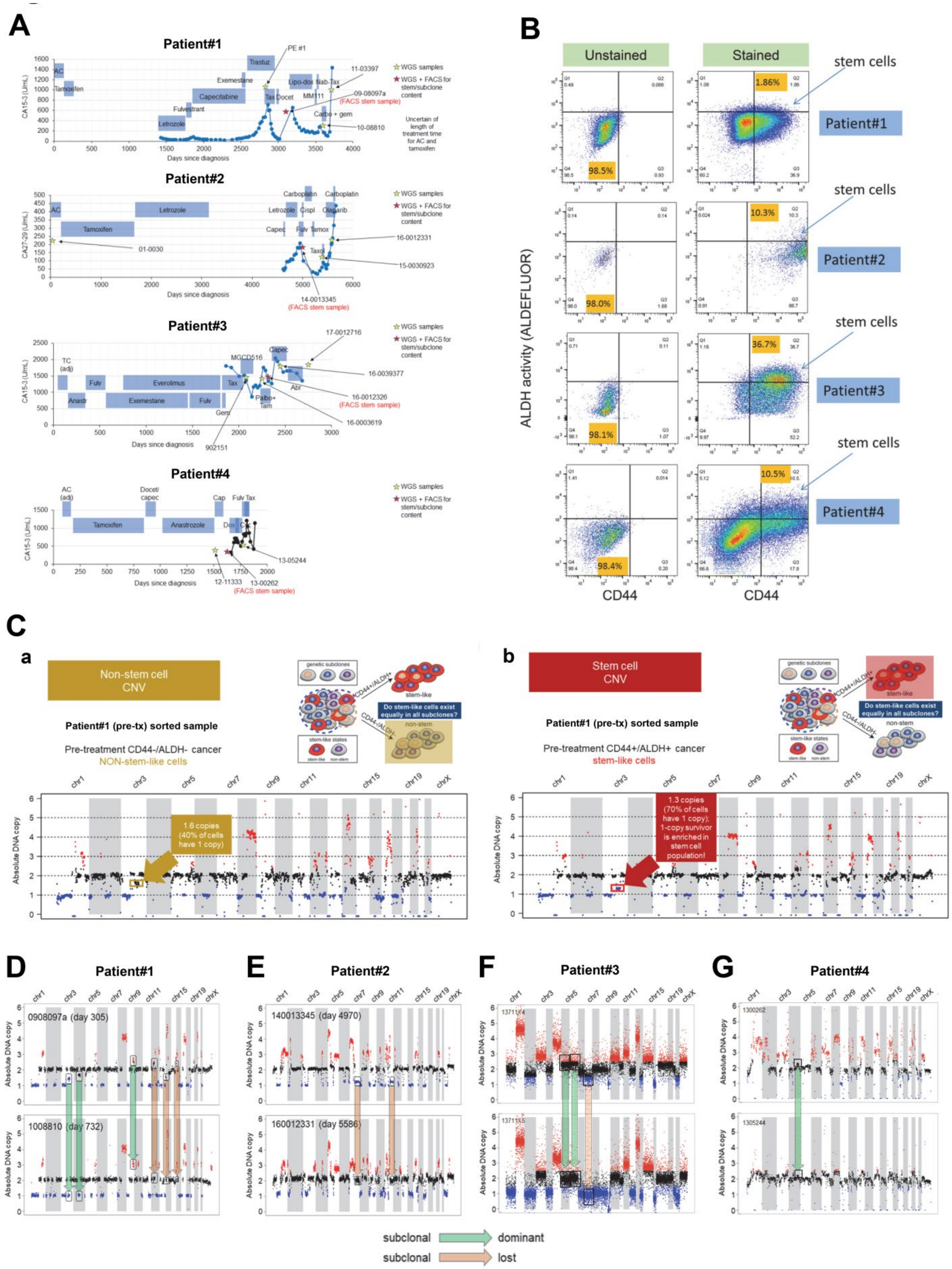
A
Patient#1
Patient#2
Patient#3
Patient#4
WGS samples
WGS + FACS for stem/subclone content
Days since diagnosis
B
Unstained
Stained
stem cells
ALDH activity (ALDEFLUOR)
CD44
C
a
Non-stem cell CNV
Patient#1 (pre-tx) sorted sample
Pre-treatment CD44-/ALDH- cancer
NON-stem-like cells
1.6 copies (40% of cells have 1 copy)
b
Stem cell CNV
Patient#1 (pre-tx) sorted sample
Pre-treatment CD44+/ALDH+ cancer
stem-like cells
1.3 copies (70% of cells have 1 copy); 1-copy survivor is enriched in stem cell population!
Do stem-like cells exist equally in all subclones?
Absolute DNA copy
D
Patient#1
0908097a (day 305)
1008810 (day 732)
E
Patient#2
140013345 (day 4970)
160012331 (day 5586)
F
Patient#3
G
Patient#4
1300262
1305244
subclonal
dominant
subclonal
lost

**Fig. S1 CNV evolution of four breast cancer patients.** (**A**) Treatment histories of four patients used for subclone/CSL state relationship analysis. (**B**) FACS sorting plots (ALDH and CD44) for negative control and untreated cells with patient pleural effusion for Patient #1-4. (**C**) Example analysis of subclone content in CSL vs. non-CSL populations. Pleural effusion 0908097a (from Patient#1) was sorted into non-CSL (**a**) or CSL (**b**) populations via ALDH/CD44 staining. DNA was then isolated and low-coverage WGS was performed. A subclonal CNV on chromosome 3 varies in prevalence in CSL cells vs. non-CSL cells. (**D-G**) Pre-treatment and post-treatment CNV analysis was performed on indicated patients using 40-60X WGS [(**D**), (**E**), and (**F**)] or low-coverage WGS (**G**). (**D**) and (**E**) have been published previously in Nature communication [19]. These two figures were allowed to be re-used due to the copyright permission of the original journal.

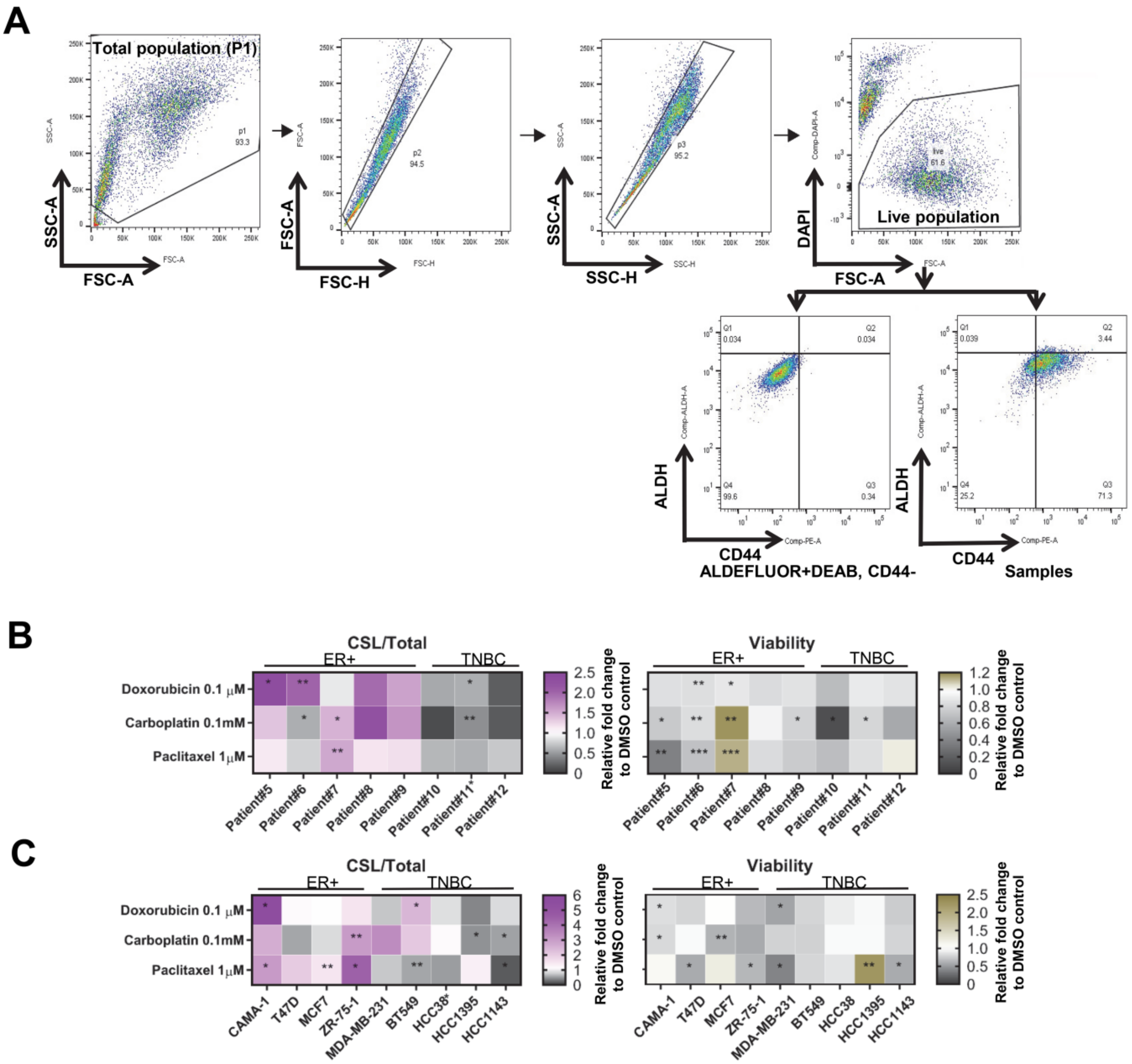
A
Total population (P1)
p1
93.3
SSC-A
FSC-A
p2
94.5
FSC-A
FSC-H
p3
95.2
SSC-A
SSC-H
live
61.6
Live population
DAPI
FSC-A
Q1
0.034
Q2
0.034
Q4
99.6
Q3
0.34
ALDH
CD44
ALDEFLUOR+DEAB, CD44-
Q1
0.039
Q2
3.44
Q4
25.2
Q3
71.3
ALDH
CD44
Samples
B
CSL/Total
ER+
TNBC
Doxorubicin 0.1 μM
Carboplatin 0.1mM
Paclitaxel 1μM
Patient#5
Patient#6
Patient#7
Patient#8
Patient#9
Patient#10
Patient#11*
Patient#12
Relative fold change to DMSO control
Viability
ER+
TNBC
Patient#5
Patient#6
Patient#7
Patient#8
Patient#9
Patient#10
Patient#11
Patient#12
Relative fold change to DMSO control
C
CSL/Total
ER+
TNBC
Doxorubicin 0.1 μM
Carboplatin 0.1mM
Paclitaxel 1μM
CAMA-1
T47D
MCF7
ZR-75-1
MDA-MB-231
BT549
HCC38*
HCC1395
HCC1143
Relative fold change to DMSO control
Viability
ER+
TNBC
CAMA-1
T47D
MCF7
ZR-75-1
MDA-MB-231
BT549
HCC38
HCC1395
HCC1143
Relative fold change to DMSO control

**Fig. S2 Chemotherapy promotion on CSL state in breast cancer.** (**A**) FACS gating strategy used to analyze breast cancer tumor cells and cell lines. (**B-C**) Heat maps for viability (live population/p1), and CSL cells/Total (ALDH+, CD44+/p1) with chemotherapy treatment in cultured patient cells (**B**) and breast cancer cell lines (**C**). The cell culture, treatment and measurement were as indicated in Fig. 2. The treatments with vehicle (DMSO) in each patient sample and cell line served as the controls and were set as fold one. All the chemotherapy treatments were expressed as the fold change relative to the controls. The mean values of triplicate tests were showed with color indicated as the legends for each treatment. Student's t-tests were performed between each treatment vs. control with triplicate tests. Significance were marked with * for $P<0.05$, ** for $P<0.01$, *** for $P<0.001$.

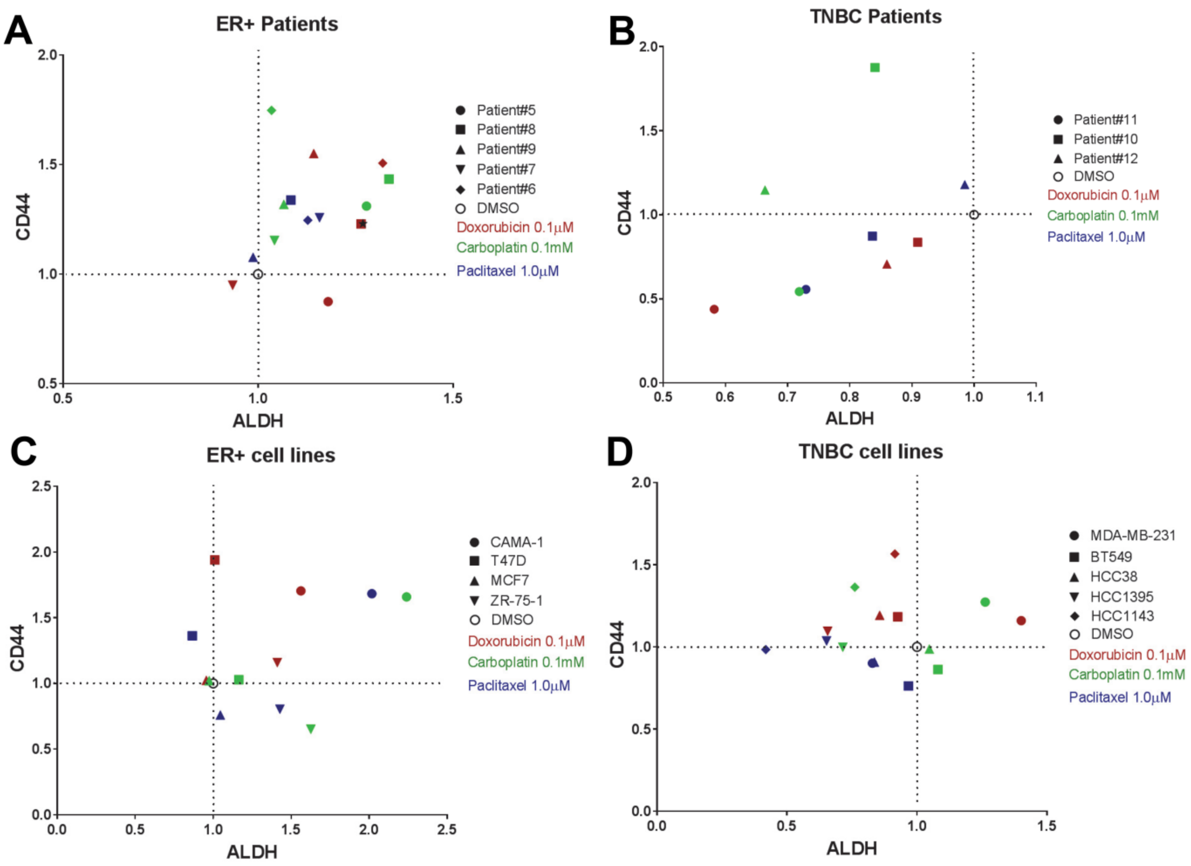
A
ER+ Patients
CD44
ALDH
Patient#5
Patient#8
Patient#9
Patient#7
Patient#6
DMSO
Doxorubicin 0.1µM
Carboplatin 0.1mM
Paclitaxel 1.0µM
B
TNBC Patients
Patient#11
Patient#10
Patient#12
C
ER+ cell lines
CAMA-1
T47D
MCF7
ZR-75-1
D
TNBC cell lines
MDA-MB-231
BT549
HCC38
HCC1395
HCC1143

**Fig. S3 ALDH-CD44 XY plots for chemotherapy promoted CSL state.** (**A-D**) Mean values of ALDH and CD44 were indicated in 2D XY plots with ER+ breast cancer patient tumor cells (**A**), TNBC breast cancer patient tumor cells (**B**), ER+ breast cancer cell lines (**C**), and TNBC breast cancer cell lines (**D**). DMSO control was marked with O (ALDH fold 1, CD44 fold 1) in each figure with broken lines connected to X and Y axis to separate the CSL promotion and reversal area. For each spot, the mean value from triplicate tests was shown for each patient sample or cell line with indicated chemotherapy treatment as shown in the legends.

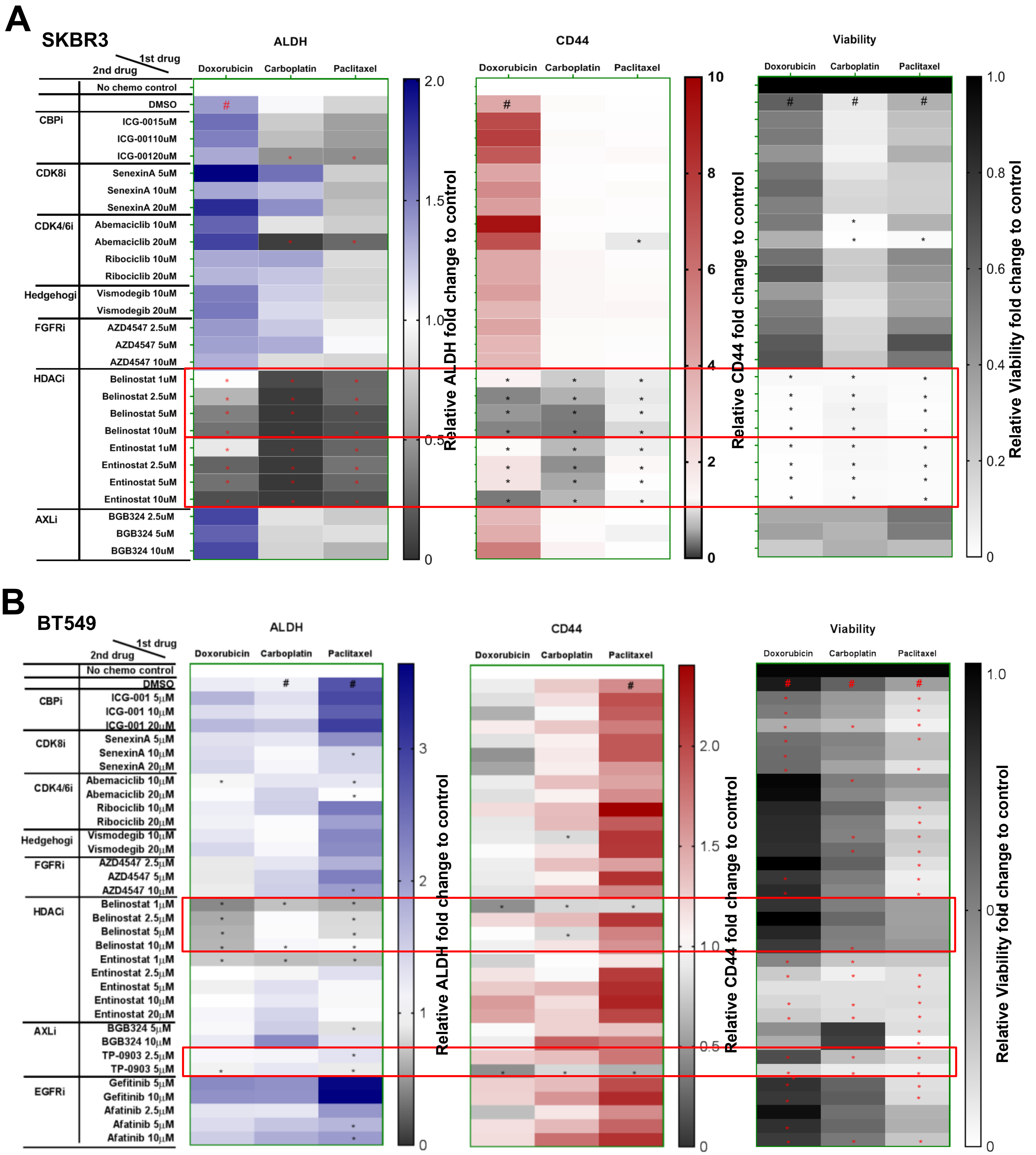
A
SKBR3
1st drug
2nd drug
ALDH
CD44
Viability
Doxorubicin
Carboplatin
Paclitaxel
No chemo control
DMSO
CBPi
ICG-0015uM
ICG-00110uM
ICG-00120uM
CDK8i
SenexinA 5uM
SenexinA 10uM
SenexinA 20uM
CDK4/6i
Abemaciclib 10uM
Abemaciclib 20uM
Ribociclib 10uM
Ribociclib 20uM
Hedgehogi
Vismodegib 10uM
Vismodegib 20uM
FGFRi
AZD4547 2.5uM
AZD4547 5uM
AZD4547 10uM
HDACi
Belinostat 1uM
Belinostat 2.5uM
Belinostat 5uM
Belinostat 10uM
Entinostat 1uM
Entinostat 2.5uM
Entinostat 5uM
Entinostat 10uM
AXLi
BGB324 2.5uM
BGB324 5uM
BGB324 10uM
Relative ALDH fold change to control
Relative CD44 fold change to control
Relative Viability fold change to control
B
BT549
ICG-001 5μM
ICG-001 10μM
ICG-001 20μM
SenexinA 5μM
SenexinA 10μM
SenexinA 20μM
Abemaciclib 10μM
Abemaciclib 20μM
Ribociclib 10μM
Ribociclib 20μM
Vismodegib 10μM
Vismodegib 20μM
AZD4547 2.5μM
AZD4547 5μM
AZD4547 10μM
Belinostat 1μM
Belinostat 2.5μM
Belinostat 5μM
Belinostat 10μM
Entinostat 1μM
Entinostat 2.5μM
Entinostat 5μM
Entinostat 10μM
Entinostat 20μM
BGB324 5μM
BGB324 10μM
TP-0903 2.5μM
TP-0903 5μM
EGFRi
Gefitinib 5μM
Gefitinib 10μM
Afatinib 2.5μM
Afatinib 5μM
Afatinib 10μM

**Fig. S4 Inhibition on chemotherapy-promoted CSL state by belinostat in additional cell lines.** Heat maps of FACS mean values for CSL reversal drugs screening with SKBR3 (**A**) and BT549 (**B**) cell lines. Cells were cultured in medium (3D) and treated for 72 hours with doxorubicin (0.1μM for SKBR3, 0.5μM for BT549), carboplatin (50μM for all cell lines), and paclitaxel (1μM for all cell lines), and then treated with potential CSL inhibitors as indicated doses for 72 hours before ALDEFLUOR/CD44 staining. The FACS mean values of ALDH and CD44 in live populations and viability were shown in the heat maps. The vehicle controls (DMSO+DMSO) were set as fold one. All the other chemotherapy plus DMSO/inhibitor combinations were expressed as the fold change relative to the controls. The Chemo plus DMSO served as the positive controls. The mean values of triplicate tests were showed with color indicated as the legends. In heat maps of ALDH and CD44, # is chemo plus DMSO significantly higher ($p<0.05$) than no treatment control, and in heat maps of viability, # is chemo plus DMSO significantly lower ($p<0.05$) than no treatment control. In all heat maps, * is chemo plus inhibitor significantly lower ($p<0.05$) than chemo plus DMSO. The potential inhibitors are highlighted with red blocks as its reversal effect for both ALDH and CD44.

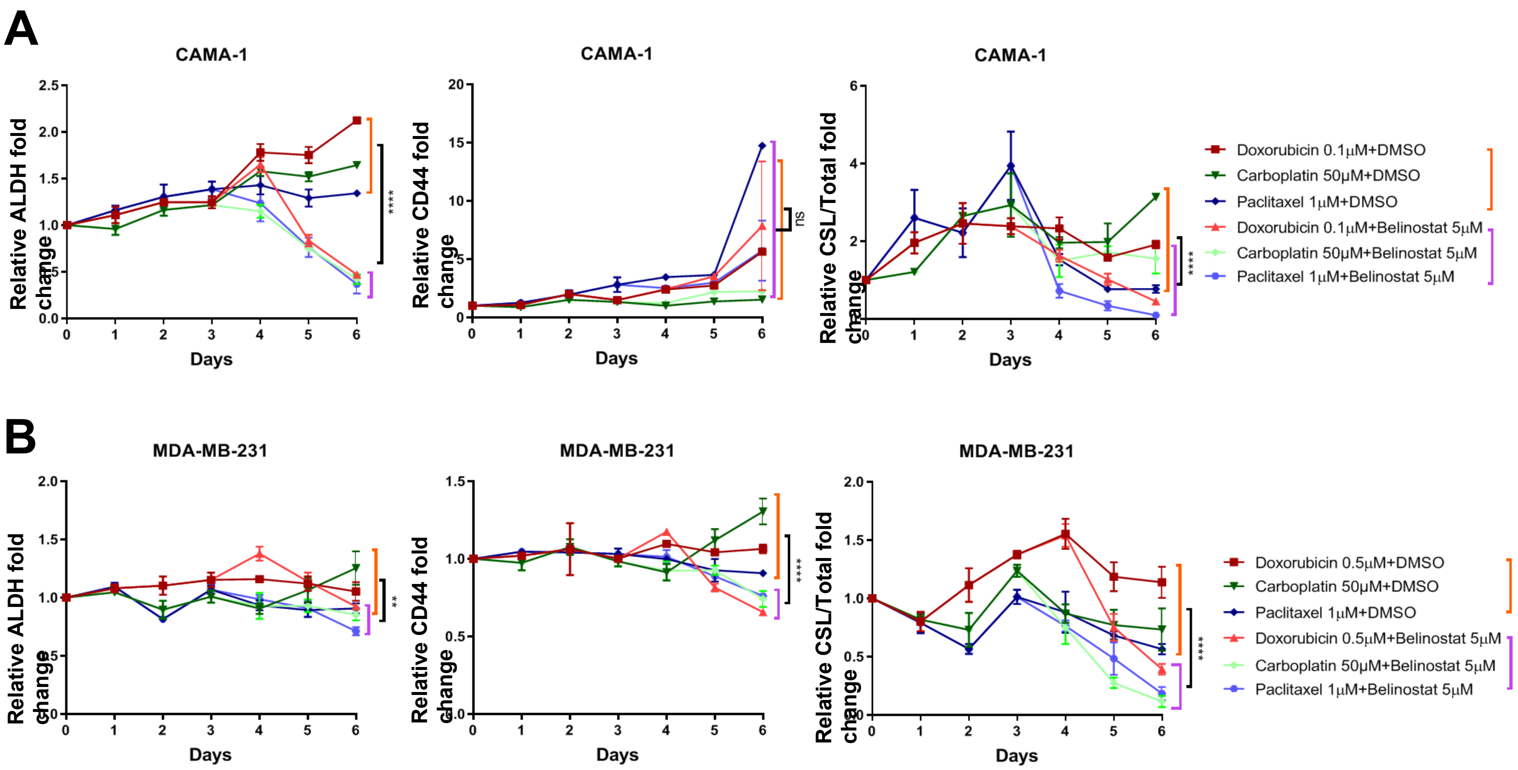
A
CAMA-1
Relative ALDH fold change
Days
****
CAMA-1
Relative CD44 fold change
Days
ns
CAMA-1
Relative CSL/Total fold change
Days
****
Doxorubicin 0.1μM+DMSO
Carboplatin 50μM+DMSO
Paclitaxel 1μM+DMSO
Doxorubicin 0.1μM+Belinostat 5μM
Carboplatin 50μM+Belinostat 5μM
Paclitaxel 1μM+Belinostat 5μM
B
MDA-MB-231
Relative ALDH fold change
Days
**
MDA-MB-231
Relative CD44 fold change
Days
****
MDA-MB-231
Relative CSL/Total fold change
Days
****
Doxorubicin 0.5μM+DMSO
Carboplatin 50μM+DMSO
Paclitaxel 1μM+DMSO
Doxorubicin 0.5μM+Belinostat 5μM
Carboplatin 50μM+Belinostat 5μM
Paclitaxel 1μM+Belinostat 5μM

**Fig. S5 Time-course FACS analysis for CAMA-1 and MDA-MB-231 with chemo plus belinostat treatment.** The FACS mean value of ALDH, CD44, and CSL cells/Total (ALDH+, CD44+/p1) were expressed with curves for CAMA-1(**A**) and MDA-MB-231 (**B**). The cell culture, treatment, measurement, and quantification were as indicated in Fig. 3B. In each day, DMSO treatment served as control and was set as fold one. All the other treatments were expressed as the fold values relative to the controls. Each spot represents the mean value of triplicate tests. The differences between chemo plus DMSO (9 replicates from three chemo plus DSMO) and chemo plus belinostat (9 replicates from three chemo plus belinostat) at Day 6 were evaluated by student's T tests. Significance is marked with * for $P<0.05$, ** for $P<0.01$, *** for $P<0.001$, *** for $P<0.0001$, **** for $P<0.0001$.

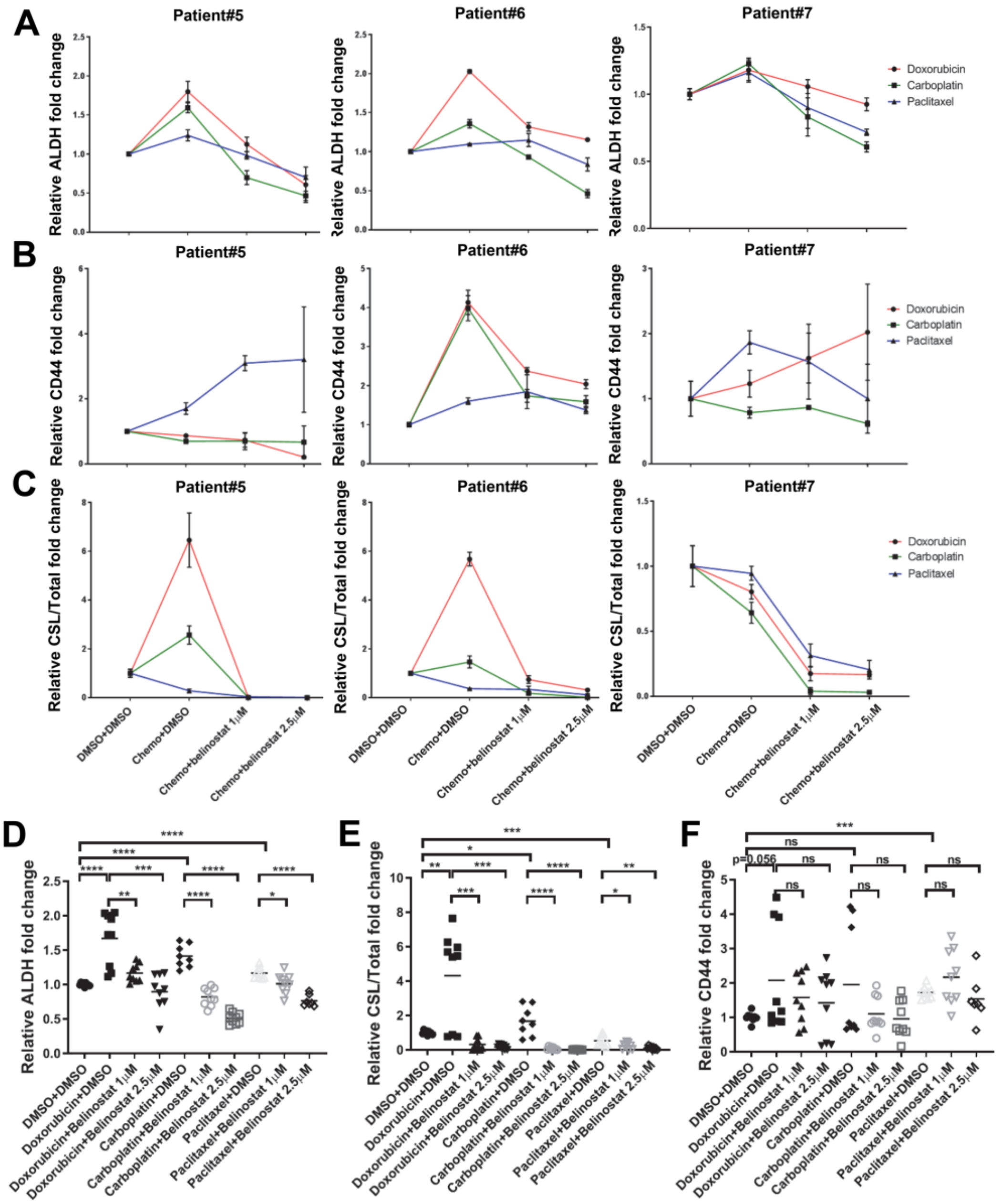
A
B
C
D
E
F
Patient#5
Patient#6
Patient#7
Relative ALDH fold change
Relative CD44 fold change
Relative CSL/Total fold change
Doxorubicin
Carboplatin
Paclitaxel
DMSO+DMSO
Chemo+DMSO
Chemo+belinostat 1μM
Chemo+belinostat 2.5μM
Doxorubicin+DMSO
Doxorubicin+Belinostat 1μM
Doxorubicin+Belinostat 2.5μM
Carboplatin+DMSO
Carboplatin+Belinostat 1μM
Carboplatin+Belinostat 2.5μM
Paclitaxel+DMSO
Paclitaxel+Belinostat 1μM
Paclitaxel+Belinostat 2.5μM
p=0.056
ns

**Fig. S6 Reversal of chemo-induced CSL state by belinostat in ER+ patient samples.** The FACS mean value of ALDH (**A**), CD44 (**B**) and CSL cells/Total (ALDH+, CD44+ cells /p1) (**C**) were expressed with curves for Patient#5, #6, and #7. The cell culture, treatment, measurement, and quantification were as indicated in Fig. 5. DMSO treatment only (DMSO+DMSO) served as controls and was set as fold one. All the other treatments were expressed as the fold values relative to the controls. Each spot in (**A-C**) represents the mean values of triplicate tests. Column graphs with individual values were generated by combining 9 replicates from three patients for each treatment to evaluate the changes in FACS mean value of ALDH (**D**), CSL cells/Total (**E**) and CD44(**F**) across patient samples. The statistical analysis was performed using student's t-tests between the indicated groups. Significance were marked with * for $P<0.05$, ** for $P<0.01$, *** for $P<0.001$, **** for $P<0.0001$.

A

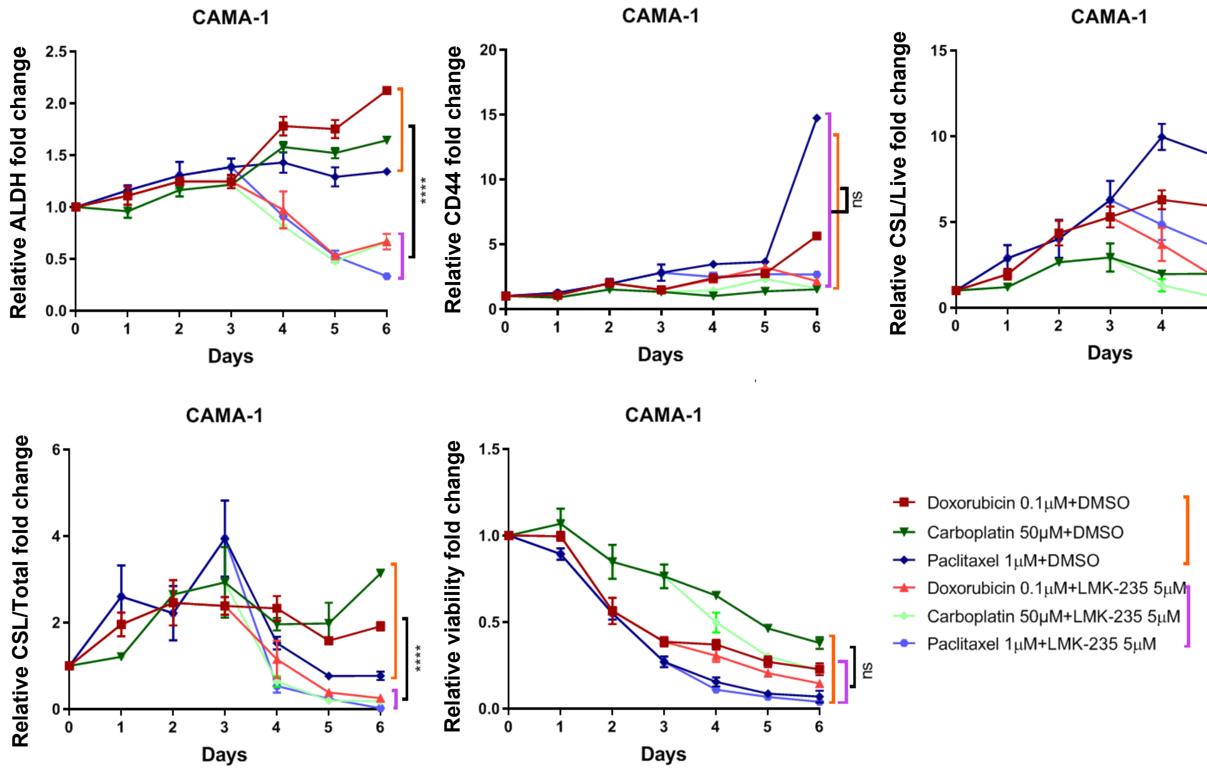
CAMA-1
Relative ALDH fold change
Days
****
CAMA-1
Relative CD44 fold change
Days
ns
CAMA-1
Relative CSL/Live fold change
Days
****
CAMA-1
Relative CSL/Total fold change
Days
****
CAMA-1
Relative viability fold change
Days
ns
Doxorubicin 0.1μM+DMSO
Carboplatin 50μM+DMSO
Paclitaxel 1μM+DMSO
Doxorubicin 0.1μM+LMK-235 5μM
Carboplatin 50μM+LMK-235 5μM
Paclitaxel 1μM+LMK-235 5μM

B

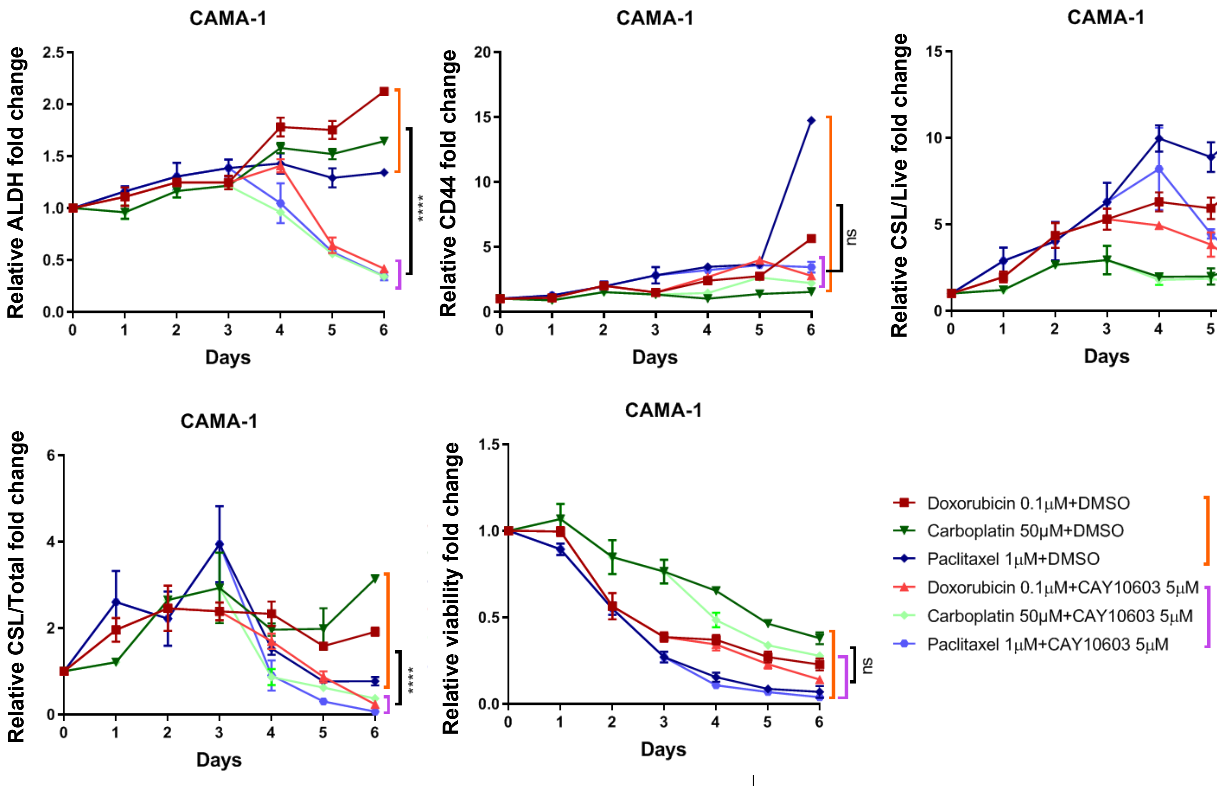
CAMA-1
Relative ALDH fold change
Days
****
CAMA-1
Relative CD44 fold change
Days
ns
CAMA-1
Relative CSL/Live fold change
Days
***
CAMA-1
Relative CSL/Total fold change
Days
****
CAMA-1
Relative viability fold change
Days
ns
Doxorubicin 0.1μM+DMSO
Carboplatin 50μM+DMSO
Paclitaxel 1μM+DMSO
Doxorubicin 0.1μM+CAY10603 5μM
Carboplatin 50μM+CAY10603 5μM
Paclitaxel 1μM+CAY10603 5μM

**Fig. S7 Reversal of chemotherapy-promoted CSL state by HDAC class II inhibitors in ER+ cells.** Line figures of relative fold changes for FACS mean value of ALDH and CD44, CSL cells/Live, CSL cells/Total and viability were shown for CAMA-1 with treatments of LMK235 (**A**) and CAY10693 (**B**). CAMA-1 cells were cultured in medium (3D) and treated for 72 hours with doxorubicin (0.1μM), carboplatin (50μM), and paclitaxel (1μM), and then replaced with medium containing LMK-235 (5μM) or CAY10603 (5μM) for 72 hours incubation. Samples were collected every day from Day 1 to Day 6, and followed by ALDEFLUOR/CD44/DAPI staining and FACS analysis. DMSO treatment only in each day served as controls and was set as fold one. All the other treatments were expressed as the fold values relative to the controls. Each spot represents the mean values of triplicate tests. The differences between chemo plus DMSO (9 replicates from three chemo plus DSMO) and chemo plus belinostat (9 replicates from three chemo plus belinostat) at Day 6 were evaluated by student's T tests. Significance is marked with * for P<0.05, ** for P<0.01, *** for P<0.001, **** for P<0.0001.

**A**

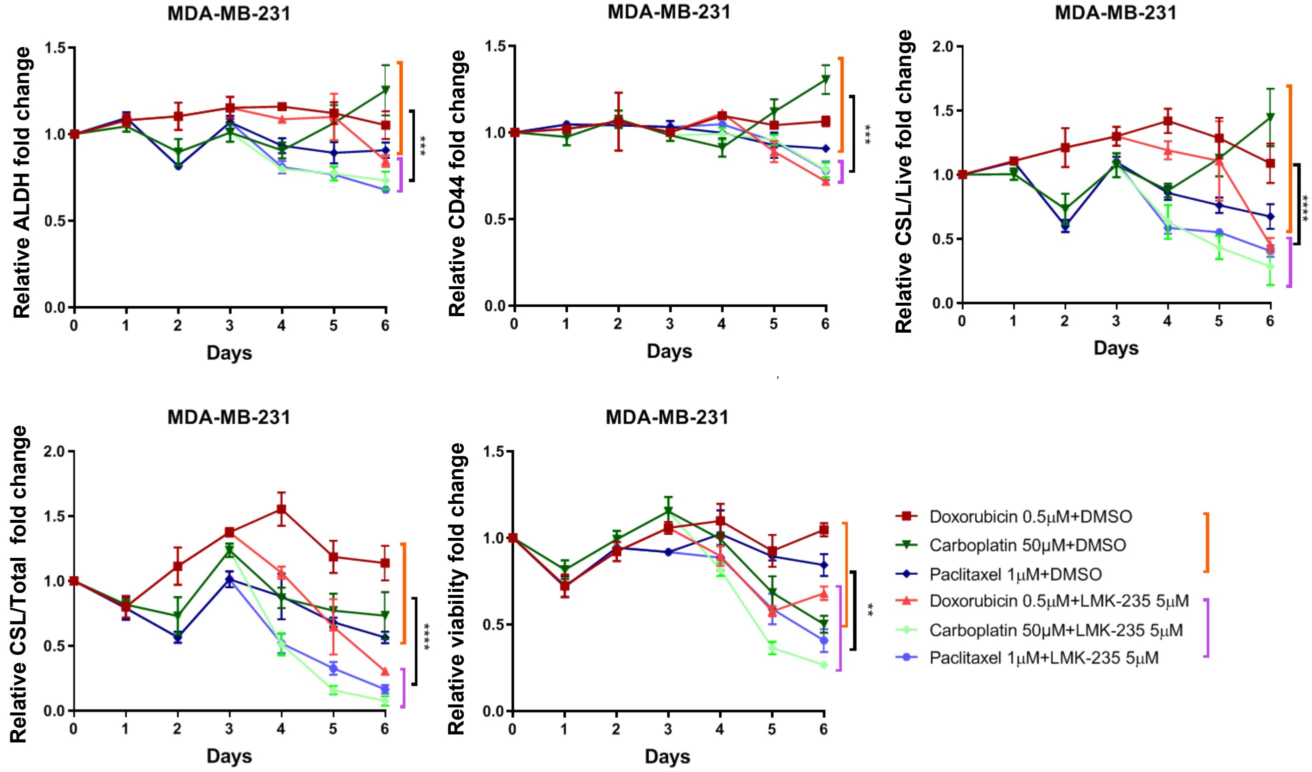
MDA-MB-231
Relative ALDH fold change
Relative CD44 fold change
Relative CSL/Live fold change
Relative CSL/Total fold change
Relative viability fold change
Days
***
***
****
****
**
Doxorubicin 0.5μM+DMSO
Carboplatin 50μM+DMSO
Paclitaxel 1μM+DMSO
Doxorubicin 0.5μM+LMK-235 5μM
Carboplatin 50μM+LMK-235 5μM
Paclitaxel 1μM+LMK-235 5μM

**B**

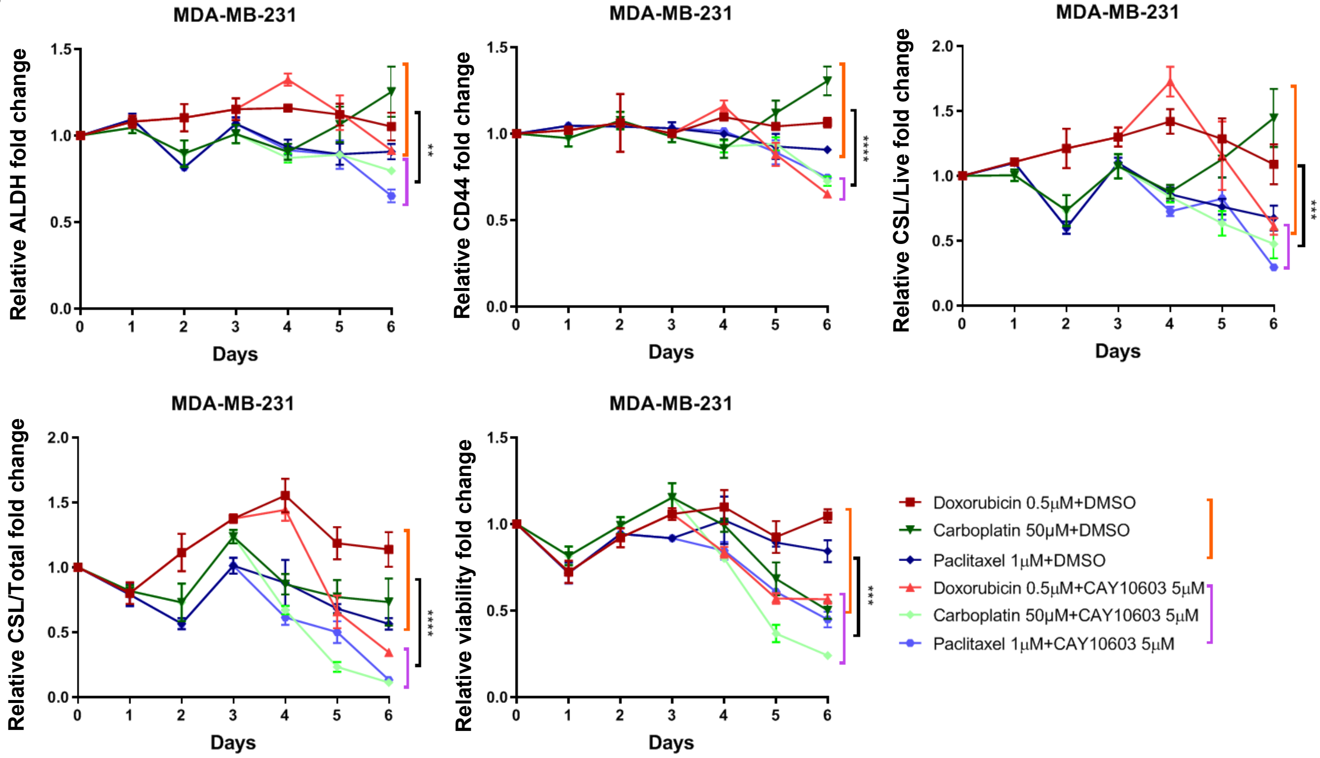
MDA-MB-231
Relative ALDH fold change
Relative CD44 fold change
Relative CSL/Live fold change
Relative CSL/Total fold change
Relative viability fold change
Days
**
****
***
****
***
Doxorubicin 0.5μM+DMSO
Carboplatin 50μM+DMSO
Paclitaxel 1μM+DMSO
Doxorubicin 0.5μM+CAY10603 5μM
Carboplatin 50μM+CAY10603 5μM
Paclitaxel 1μM+CAY10603 5μM

**Fig. S8 Reversal of chemotherapy-promoted CSL state by HDAC class II inhibitors in TNBC cells.** Line figures of relative fold changes for FACS mean value of ALDH and CD44, CSL cells/Live, CSL cells/Total and viability were shown for MDA-MB-231 with treatments of LMK235 (**A**) and CAY10693 (**B**). MDA-MB-231 cells were cultured in medium (3D) and treated for 72 hours with doxorubicin (0.5μM), carboplatin (50μM), and paclitaxel (1μM), and then replaced with medium containing LMK-235 (5μM) or CAY10603 (5μM) for 72 hours incubation. Samples were collected every day from Day 1 to Day 6, and followed by ALDEFLUOR/CD44/DAPI staining and FACS analysis. DMSO treatment only in each day served as controls and was set as fold one. All the other treatments were expressed as the fold values relative to the controls. Each spot represents the mean values of triplicate tests. The differences between chemo plus DMSO (9 replicates from three chemo plus DSMO) and chemo plus belinostat (9 replicates from three chemo plus belinostat) at Day 6 were evaluated by student's T tests. Significance is marked with * for P<0.05, ** for P<0.01, *** for P<0.001, **** for P<0.0001.

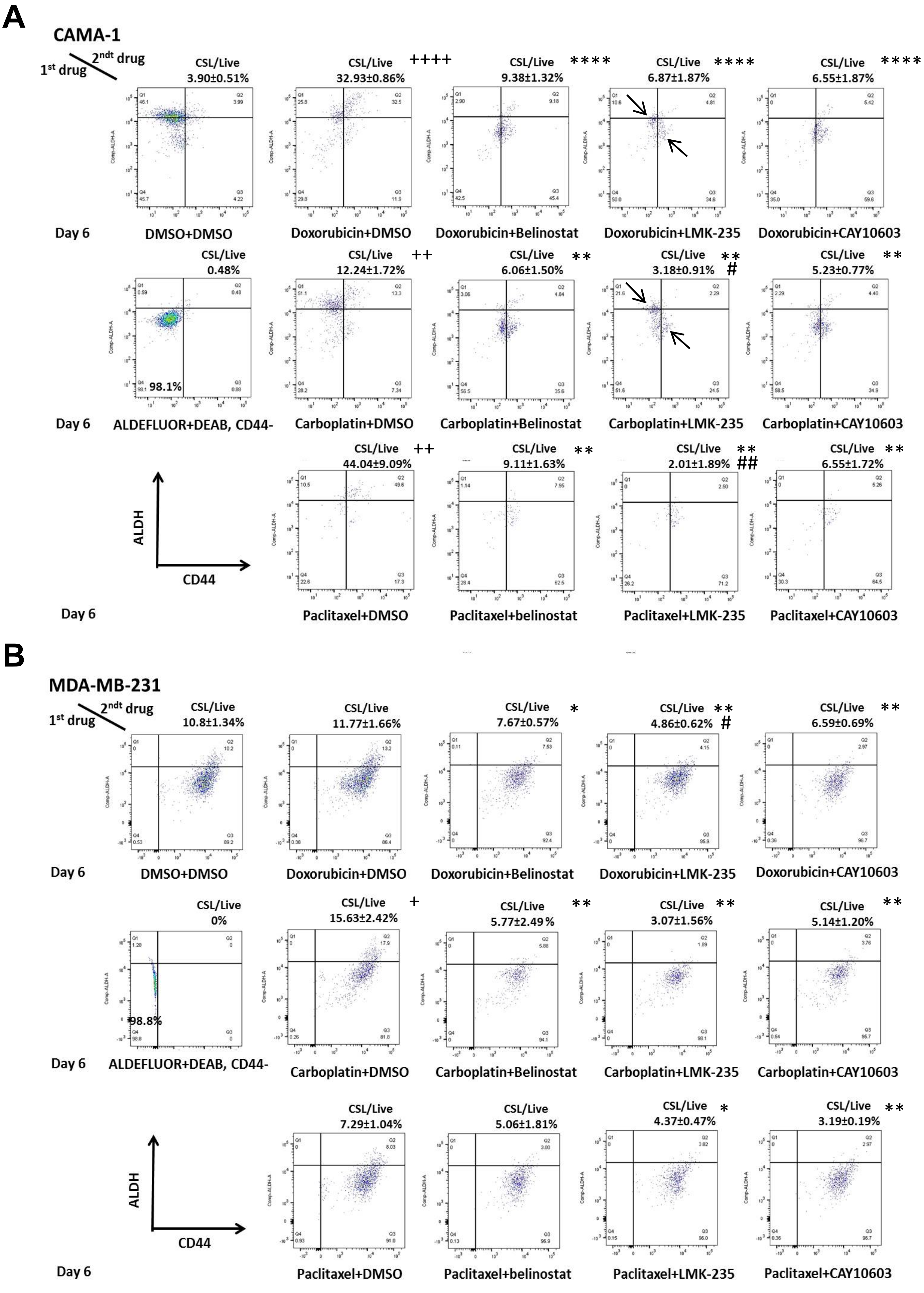
A
CAMA-1
1st drug
2ndt drug
CSL/Live
3.90±0.51%
DMSO+DMSO
CSL/Live ++++
32.93±0.86%
Doxorubicin+DMSO
CSL/Live ****
9.38±1.32%
Doxorubicin+Belinostat
CSL/Live ****
6.87±1.87%
Doxorubicin+LMK-235
CSL/Live ****
6.55±1.87%
Doxorubicin+CAY10603
Day 6
CSL/Live
0.48%
98.1%
ALDEFLUOR+DEAB, CD44-
CSL/Live ++
12.24±1.72%
Carboplatin+DMSO
CSL/Live **
6.06±1.50%
Carboplatin+Belinostat
CSL/Live **
3.18±0.91% #
Carboplatin+LMK-235
CSL/Live **
5.23±0.77%
Carboplatin+CAY10603
Day 6
ALDH
CD44
CSL/Live ++
44.04±9.09%
Paclitaxel+DMSO
CSL/Live **
9.11±1.63%
Paclitaxel+belinostat
CSL/Live **
2.01±1.89% ##
Paclitaxel+LMK-235
CSL/Live **
6.55±1.72%
Paclitaxel+CAY10603
Day 6
B
MDA-MB-231
1st drug
2ndt drug
CSL/Live
10.8±1.34%
DMSO+DMSO
CSL/Live
11.77±1.66%
Doxorubicin+DMSO
CSL/Live *
7.67±0.57%
Doxorubicin+Belinostat
CSL/Live **
4.86±0.62% #
Doxorubicin+LMK-235
CSL/Live **
6.59±0.69%
Doxorubicin+CAY10603
Day 6
CSL/Live
0%
98.8%
ALDEFLUOR+DEAB, CD44-
CSL/Live +
15.63±2.42%
Carboplatin+DMSO
CSL/Live **
5.77±2.49%
Carboplatin+Belinostat
CSL/Live **
3.07±1.56%
Carboplatin+LMK-235
CSL/Live **
5.14±1.20%
Carboplatin+CAY10603
Day 6
ALDH
CD44
CSL/Live
7.29±1.04%
Paclitaxel+DMSO
CSL/Live
5.06±1.81%
Paclitaxel+belinostat
CSL/Live *
4.37±0.47%
Paclitaxel+LMK-235
CSL/Live **
3.19±0.19%
Paclitaxel+CAY10603
Day 6

**Fig. S9 Detailed FACS analysis of HDAC inhibitors on chemotherapy-promoted CSL state.** Detailed time course FACS plots of ALDH vs.CD44 at Day6 were shown for live cells with chemo plus DSMO/belinostat/LMK-235/CAY10603 treatments in CAMA-1 (**A**) and MDA-MB-231(**B**), respectively. ALDH/CD44 negative controls with DEAB were used to determine the gates for CSL cells. For each treatment, the mean values from triplicate tests plus standard deviation errors were showed on the right side above each plot for the ratio of CSL/Live, The significance between each treatments were evaluated by student's T tests. + is chemo plus DMSO significantly higher ($p<0.05$) than no treatment control, * is chemo plus HDACi significantly lower ($p<0.05$) than chemo plus DMSO, and # is chemo plus soecific HDACi (LMK-235 or CAY10603) significantly lower ($p<0.05$) than chemo plus belinostat.

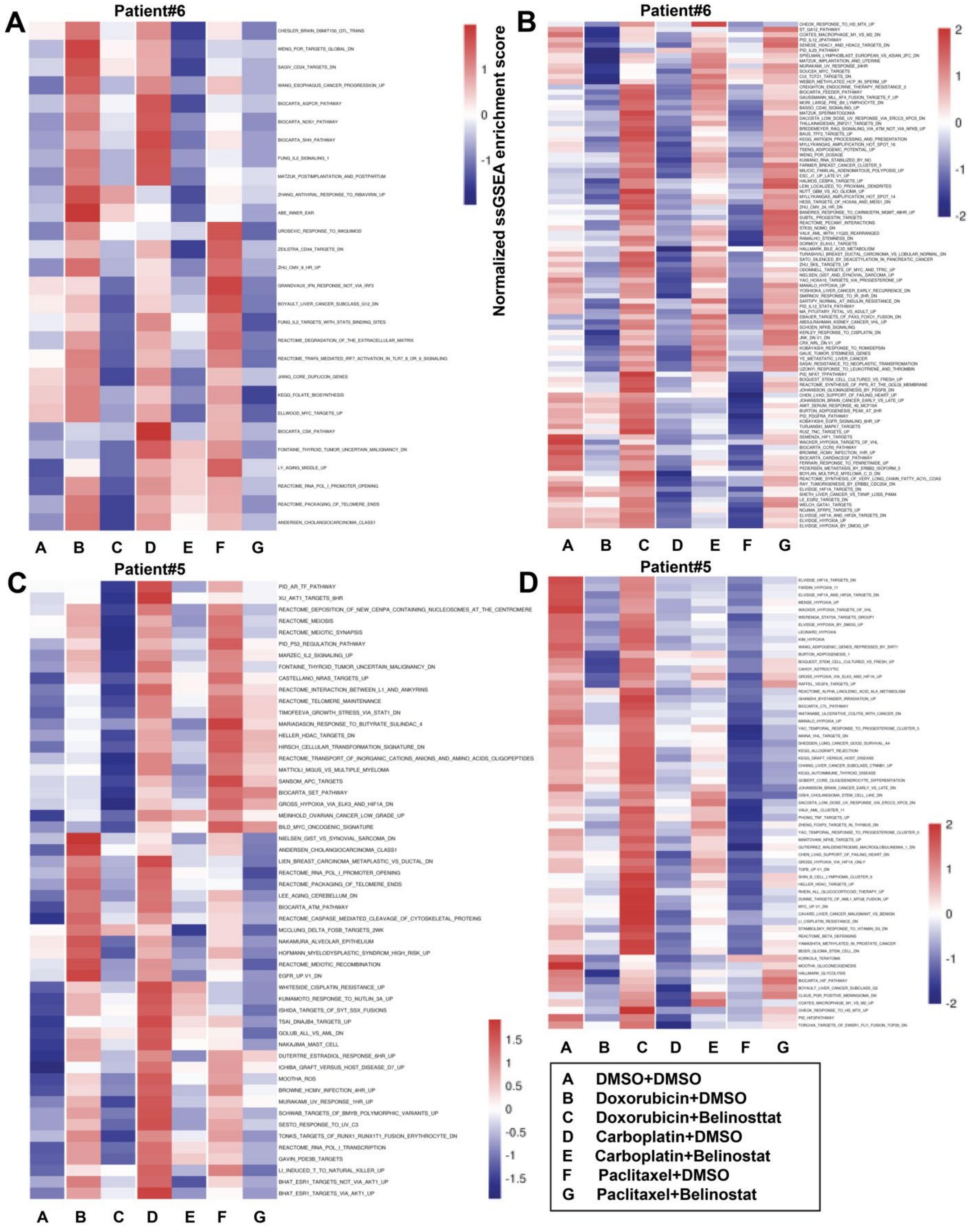
A
Patient#6
CHESLER_BRAIN_D6MIT150_QTL_TRANS
WENG_POR_TARGETS_GLOBAL_DN
SAGIV_CD24_TARGETS_DN
WANG_ESOPHAGUS_CANCER_PROGRESSION_UP
BIOCARTA_AGPCR_PATHWAY
BIOCARTA_NOS1_PATHWAY
BIOCARTA_SHH_PATHWAY
FUNG_IL2_SIGNALING_1
MATZUK_POSTIMPLANTATION_AND_POSTPARTUM
ZHANG_ANTIVIRAL_RESPONSE_TO_RIBAVIRIN_UP
ABE_INNER_EAR
UROSEVIC_RESPONSE_TO_IMIQUIMOD
ZEILSTRA_CD44_TARGETS_DN
ZHU_CMV_8_HR_UP
GRANDVAUX_IFN_RESPONSE_NOT_VIA_IRF3
BOYAULT_LIVER_CANCER_SUBCLASS_G12_DN
FUNG_IL2_TARGETS_WITH_STAT5_BINDING_SITES
REACTOME_DEGRADATION_OF_THE_EXTRACELLULAR_MATRIX
REACTOME_TRAF6_MEDIATED_IRF7_ACTIVATION_IN_TLR7_8_OR_9_SIGNALING
JIANG_CORE_DUPLICON_GENES
KEGG_FOLATE_BIOSYNTHESIS
ELLWOOD_MYC_TARGETS_UP
BIOCARTA_CSK_PATHWAY
FONTAINE_THYROID_TUMOR_UNCERTAIN_MALIGNANCY_DN
LY_AGING_MIDDLE_UP
REACTOME_RNA_POL_I_PROMOTER_OPENING
REACTOME_PACKAGING_OF_TELOMERE_ENDS
ANDERSEN_CHOLANGIOCARCINOMA_CLASS1
A B C D E F G
1
0
-1
Normalized ssGSEA enrichment score
B
Patient#6
2
1
0
-1
-2
A B C D E F G
C
Patient#5
PID_AR_TF_PATHWAY
XU_AKT1_TARGETS_6HR
REACTOME_DEPOSITION_OF_NEW_CENPA_CONTAINING_NUCLEOSOMES_AT_THE_CENTROMERE
REACTOME_MEIOSIS
REACTOME_MEIOTIC_SYNAPSIS
PID_P53_REGULATION_PATHWAY
MARZEC_IL2_SIGNALING_UP
FONTAINE_THYROID_TUMOR_UNCERTAIN_MALIGNANCY_DN
CASTELLANO_NRAS_TARGETS_UP
REACTOME_INTERACTION_BETWEEN_L1_AND_ANKYRINS
REACTOME_TELOMERE_MAINTENANCE
TIMOFEEVA_GROWTH_STRESS_VIA_STAT1_DN
MARIADASON_RESPONSE_TO_BUTYRATE_SULINDAC_4
HELLER_HDAC_TARGETS_DN
HIRSCH_CELLULAR_TRANSFORMATION_SIGNATURE_DN
REACTOME_TRANSPORT_OF_INORGANIC_CATIONS_ANIONS_AND_AMINO_ACIDS_OLIGOPEPTIDES
MATTIOLI_MGUS_VS_MULTIPLE_MYELOMA
SANSOM_APC_TARGETS
BIOCARTA_SET_PATHWAY
GROSS_HYPOXIA_VIA_ELK3_AND_HIF1A_DN
MEINHOLD_OVARIAN_CANCER_LOW_GRADE_UP
BILD_MYC_ONCOGENIC_SIGNATURE
NIELSEN_GIST_VS_SYNOVIAL_SARCOMA_DN
ANDERSEN_CHOLANGIOCARCINOMA_CLASS1
LIEN_BREAST_CARCINOMA_METAPLASTIC_VS_DUCTAL_DN
REACTOME_RNA_POL_I_PROMOTER_OPENING
REACTOME_PACKAGING_OF_TELOMERE_ENDS
LEE_AGING_CEREBELLUM_DN
BIOCARTA_ATM_PATHWAY
REACTOME_CASPASE_MEDIATED_CLEAVAGE_OF_CYTOSKELETAL_PROTEINS
MCCLUNG_DELTA_FOSB_TARGETS_2WK
NAKAMURA_ALVEOLAR_EPITHELIUM
HOFMANN_MYELODYSPLASTIC_SYNDROM_HIGH_RISK_UP
REACTOME_MEIOTIC_RECOMBINATION
EGFR_UP.V1_DN
WHITESIDE_CISPLATIN_RESISTANCE_UP
KUMAMOTO_RESPONSE_TO_NUTLIN_3A_UP
ISHIDA_TARGETS_OF_SYT_SSX_FUSIONS
TSAI_DNAJB4_TARGETS_UP
GOLUB_ALL_VS_AML_DN
NAKAJIMA_MAST_CELL
DUTERTRE_ESTRADIOL_RESPONSE_6HR_UP
ICHIBA_GRAFT_VERSUS_HOST_DISEASE_D7_UP
MOOTHA_ROS
BROWNE_HCMV_INFECTION_4HR_UP
MURAKAMI_UV_RESPONSE_1HR_UP
SCHWAB_TARGETS_OF_BMYB_POLYMORPHIC_VARIANTS_UP
SESTO_RESPONSE_TO_UV_C3
TONKS_TARGETS_OF_RUNX1_RUNX1T1_FUSION_ERYTHROCYTE_DN
REACTOME_RNA_POL_I_TRANSCRIPTION
GAVIN_PDE3B_TARGETS
LI_INDUCED_T_TO_NATURAL_KILLER_UP
BHAT_ESR1_TARGETS_NOT_VIA_AKT1_UP
BHAT_ESR1_TARGETS_VIA_AKT1_UP
1.5
1
0.5
0
-0.5
-1
-1.5
A B C D E F G
D
Patient#5
2
1
0
-1
-2
A B C D E F G
A DMSO+DMSO
B Doxorubicin+DMSO
C Doxorubicin+Belinosttat
D Carboplatin+DMSO
E Carboplatin+Belinostat
F Paclitaxel+DMSO
G Paclitaxel+Belinostat

**Fig. S10. Heat maps of ssGSEA enrichment scores for chemotherapy-belinostat reversal pathways.** Heat maps were generated based on the averages of ssGSEA enrichment scores for the shared reversal pathways. For Patient#6, Pathways increased by chemotherapy, decreased by belinostat were shown in (**A**); pathways decreased by chemotherapy, increased by belinostat were shown in (**B**). For Patient#5, pathways increased by chemotherapy, decreased by belinostat were shown in (**C**); pathways decreased by chemotherapy, increased by belinostat were shown in (**D**). Each ssGSEA enrichment scores was normalized according to the whole heat map, and expressed as the color indicated as the legends.
